# Hsa-miR-31-5p controls a metabolic switch in psoriatic keratinocytes that identifies therapeutic intervention

**DOI:** 10.1101/2022.01.21.477183

**Authors:** Mao-Jie Wang, Yong-Yue Xu, HarmJan Vos, Can Gulersonmez, Edwin Stigter, Johan Gerritsen, Marc Pages Gallego, Robert van Es, Li Li, Hao Deng, Ling Han, Run-Yue Huang, Chuan-Jian Lu, Boudewijn MT Burgering

**Affiliations:** The Second Affiliated Hospital, Guangzhou University of Chinese Medicine (Guangdong Provincial Hospital of Chinese Medicine), Guangzhou, China; Oncode Institute and Molecular Cancer Research, Center for Molecular Medicine, University Medical Center Utrecht, Utrecht, The Netherlands; Guangdong Provincial Key Laboratory of Clinical Research on Traditional Chinese Medicine Syndrome, Guangzhou, China; Metabolic Diagnostics, Department of Biomedical Genetics, Centre for Molecular Medicine, University Medical Centre Utrecht, Utrecht, The Netherlands; Oncode Institute and Department of Genetics, Center for Molecular Medicine, University Medical Center Utrecht, Utrecht, The Netherlands

## Abstract

Psoriasis is characterized by a combination of keratinocyte hyperproliferation and immune cell activation. Immune cell activation requires increased glucose consumption, consequently limiting glucose availability for other cell types like keratinocytes. In psoriasis Hsa-microRNA-31-5p (miR-31) is highly expressed in keratinocytes. Here we show that miR-31 expression in keratinocytes is induced by limited glucose availability and increases survival under limiting glucose conditions, by increasing glutamine metabolism. In addition, miR-31 induced glutamine metabolism results in secretion of specific metabolites (aspartate and glutamate) but also immuno-modulatory factors. We show that this miR-31-induced secretory phenotype is sufficient to induce Th17 cell differentiation, a hallmark of psoriasis. Inhibition of glutaminase (GLS) using CB-839 impedes miR31-induced metabolic rewiring and secretion of immuno-modulatory factors. Concordantly, pharmacological targeting of GLS alleviated psoriasis pathology in a mouse model of psoriasis. Together our data illustrate an emerging concept of metabolic interaction across cell compartments that characterizes disease development, which can be employed to design effective treatment options for disease, as shown here for psoriasis.

## Introduction

Psoriasis is a chronic immune-mediated disorder of the skin that affects 0.1-0.4% of the total world population and is typically characterized by scaly, erythematous plaques on the extensor surfaces (Armstrong & Read, 2020; Parisi *et al*, 2020). The disease causes a negative impact on quality of life and is associated with additional pathologies, such as arthritis, cardiovascular disease and metabolic syndrome (Parisi *et al*., 2020). The pathogenesis is characterized by a combination of keratinocyte hyperproliferation and inflammation mediated by activated immune cells, including neutrophils, dendritic cells and Th17 cells (Greb *et al*, 2016).

Human disease, like psoriasis, is prevented by intrinsic and extrinsic control mechanisms that will remove aberrant cells before these can survive and establish a stable altered cellular compartment. To evade these control mechanisms aberrant cells may adopt various strategies. A well-studied example being the various mechanisms by which cancer cells prevent clearance by the immune system. Immune cell activation oftentimes requires a shift in T-cell metabolism towards glycolysis and hence increased glucose consumption. Cancer cells are similarly dependent on increased glucose consumption to sustain proliferation (Warburg effect) and cancer cells have been suggested to outcompete immune cells for glucose availability and thereby establish immune suppression(Garcia-Bermudez *et al*, 2018).

Importantly, in inflammatory skin diseases like psoriasis the activated T-cells display a high glycolytic demand and consequently in these diseases T-cells may likely outcompete other cell types of glucose. However, in certain skin diseases inflammation co-occurs with increased proliferation of skin cells, mostly keratinocytes. Apparently, keratinocytes have to adopt a metabolic switch that enables them to proliferate without competing for the metabolic resources that are essential to fuel immune cell differentiation and activation resulting in inflammation. Cellular adaptation oftentimes requires changes in gene expression programs and these can be brought about through epigenetic changes, but also by expression of microRNAs (miRNAs), noncoding RNAs of 21–25 nucleotides that can regulate mRNA and consequent protein expression of hundreds of genes simultaneously.

A number of miRNAs have been implicated in psoriasis pathology (Joyce *et al*, 2011), including microRNA-31-5p (miR-31(Wang *et al*, 2019; Xu *et al*, 2013)). Interestingly, miR-31 expression is also implicated in various types of cancer (reviewed in (Laurila & Kallioniemi, 2013; Valastyan & Weinberg, 2010)) and appears therefore associated in general with increased proliferation. Therefore, we chose to study the role of miR-31 in psoriasis pathology and in order to obtain a more integrated view on miR-31 function and regulation of keratinocyte proliferation, we used a combination of proteomics and metabolomics and focused on the regulation of metabolic processes by miR-31.

Interestingly, we find that miR-31 expression impacts on glutamine metabolism at several levels. Glutamine through reductive or oxidative metabolism can provide cells with the majority of building blocks and efficiently maintain cell viability even in the absence of glucose (Altman *et al*, 2016). In agreement, miR-31 expression in keratinocytes enables survival and proliferation of keratinocytes under limiting glucose conditions by switching to glutamine dependent metabolism. Interestingly, miR-31 expression itself is induced by limiting glucose availability, suggesting that increased glucose consumption by T-cells triggers miR-31 expression in keratinocytes in psoriasis. This reciprocal interaction is further enforced by our observation that miR-31 expression results in glutamine-dependent secretion of metabolites and cytokines that enable Th17 differentiation. Consequently, breaking the cross-talk between T-cells and keratinocytes by inhibiting glutaminase (GLS) a key enzyme in glutamine metabolism, alleviates psoriasis in a mouse model. Taken together our results illustrate how the extracellular environment can be involved in disease progression and how this knowledge may be employed to tailor treatment.

## Results

First, we confirmed increased miR-31 expression in psoriasis by comparing skin biopsies from healthy control and psoriatic patients (Fig. 1a). We observed similar increase in miR-31 expression in publicly available data (Extended data Fig. 1a) and in the imiquimod (IMQ)-induced mouse model for psoriasiform hyperplasia(van der Fits *et al*, 2009) (Fig. 1a). Potential target mRNAs have been identified for miR-31 (reviewed in (Laurila & Kallioniemi, 2013)), yet mostly these proposed targets have been studied on an individual basis, whereas a specific miRNA can target simultaneously up to a few hundred of different mRNA species and thus a simultaneous change in the expression of many proteins. Importantly, this indicates that miRNAs regulate biology in a systemic rather than singular manner. To study the full spectrum of miRNA-31-induced protein deregulation we made use of a quantitative proteomic strategy (SILAC, Extended data Fig. 1b). Ectopic miR-31 expression in two different human cell lines, 293T and the keratinocyte cell line HaCAT, induced extensive changes in protein levels (Extended data Fig. 1c). Pearson correlation indicated a high concordance between SILAC proteome results obtained with these cell lines (R2=0.449, p<0.001, Extended data Fig. 1d) and a significant overlap in both cell lines of down regulated (279) and upregulated (187) proteins (Fig. 1b). To further evaluate the quality of our proteomics results we selected a set of validated miR-31 targets (meaning a target that at least is validated by reporter assay, western blot, or qPCR according miRTarBase database (http://mirtarbase.mbc.nctu.edu.tw/), Extended table 1). Comparison with our results confirmed regulation of several of these validated miR-31 targets in both cell lines (e.g., Hypoxia Inducing Factor1 alpha inhibitor (HIFAN), Forkhead box-O transcription factor 3 (FOXO3), NUMB-endocytic adaptor protein (NUMB), AT-rich interaction domain 1A (ARID1A) (Extended data Fig. 1e). Next, we checked our data for expression of 301 metabolic genes and 137 of these were detected in all replicates. When compiling these data, we identified 15 high confidence hits for metabolic proteins downregulated following miR-31 expression. Importantly, a majority of downregulated metabolic genes (13/15) harbour at least one miR-31 seed sequence (Fig. 1c, Extended table 2).

**Figure 1.**
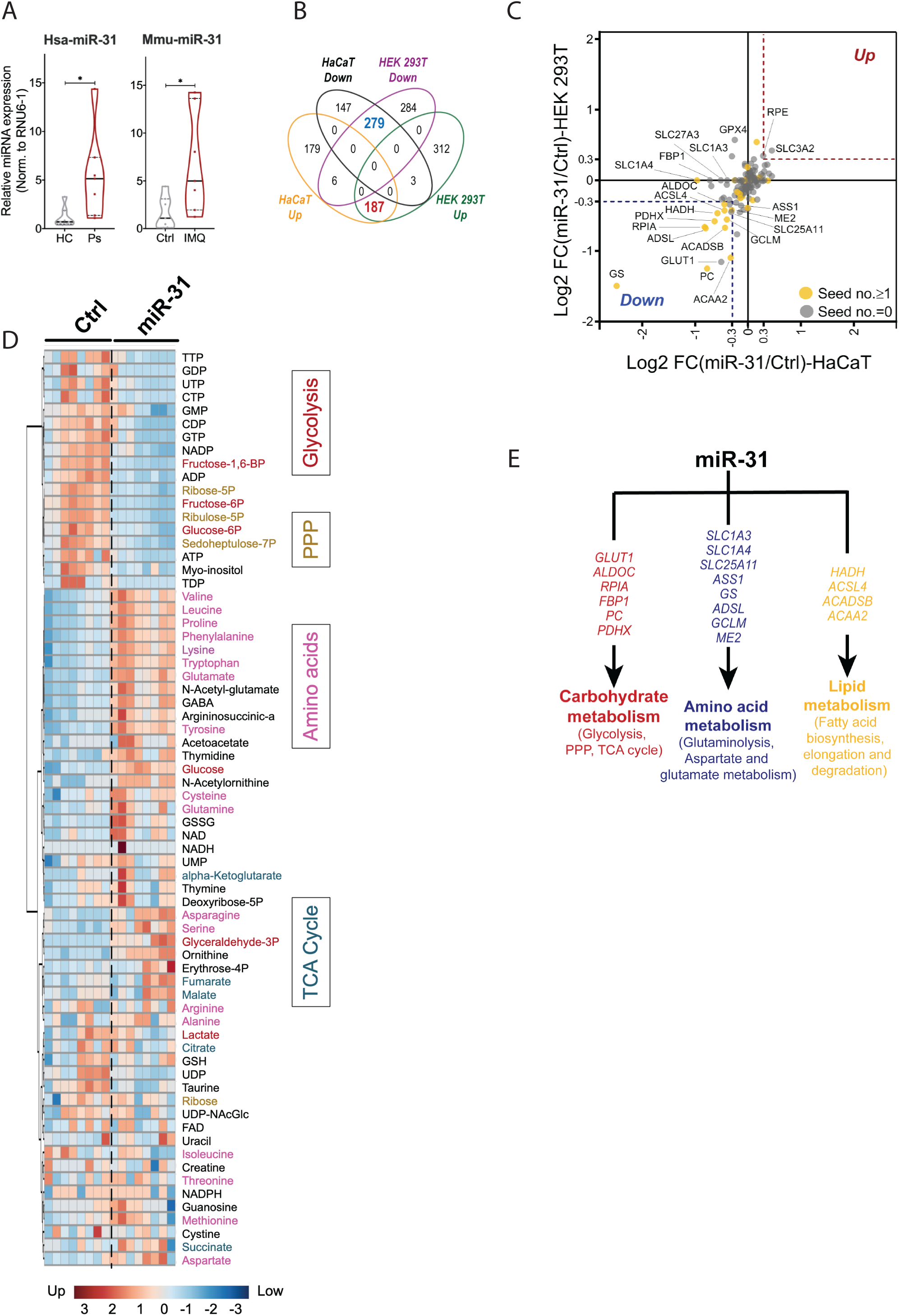
**A. miR-31 is upregulated in human psoriasis and the IMQ mouse model**. Violin plot of qRT-PCR for miR-31-5p in skin biopsies of healthy control people (HC), psoriasis patient (Ps), control mouse (Ctrl), and imiquimod-induced psoriatic mouse model (IMQ). RUN6-1 was used as house control. n=7, * means p<0.05, which was calculated by Student’s t-test. **B. miR-31 regulation of protein expression**. The Venn diagram depicts the number proteins that were either up- or down-regulated in HaCaT cells and HEK 293T cells after miR-31 expression. **C. Metabolic enzymes regulated by miR-31**. Dot plot representation of log2 fold changes of metabolic gene expression in HEK 293T cell (y axis) and HaCaT cell (x axis). Significantly downregulated or upregulated genes in both cell lines are circled by blue or red dashed outlines, respectively. Yellow marked genes contain at least one binding seed in 3’UTR for miR-31-5p. **D. Relative metabolite levels after miR-31 expression**. Heat map representation of metabolite profile in HEK 293T cells upon miR-31 overexpression. Metabolites are grouped and marked with various colours for different metabolic pathways. **E.** Summary of the impact of miR-31 on cell metabolism.

As miR-31 expression resulted in significant metabolic protein expression changes we next performed targeted analysis of the cellular metabolome using Hydrophilic Interaction Chromatography - Mass Spectrometry (HILIC-MS) to obtain an unbiased view on metabolic changes induced by miR-31 expression. Principal component analysis (PCA, Extended data Fig. 1f) and heatmap clustering (Fig. 1d, Extended table 3) showed a clear separation in metabolite expression profiles between HaCAT cells expressing miR-31 or a scrambled miRNA mimic. Importantly, we observed metabolite changes to corroborate protein expression changes. For example, in agreement with downregulation of ribose 5-phosphate isomerase A (RPIA) at the protein level and its role in the Pentose Phosphate Pathway (PPP), we observed a reduced level of PPP intermediates (Ribulose-5P, Glucose-6P, Seduheptulose-7P). Also changes in glutamine/glutamate levels are consistent with the observed downregulation of Glutamine synthetase (GS) in the proteome analysis. Pathway GO-term analysis using metabolite levels as input also indicated extensive miR-31 induced deregulation of glucose and glutamine metabolism (Extended data Fig. 1g) Combined the proteomic and metabolic data indicate that miR-31 expression, through simultaneous deregulation of the expression of multiple metabolic genes, may impact glycolytic, lipid and amino acid metabolism (summarized in Fig. 1e) and deregulated gene expression appears to correlate by and large with changes in metabolite levels.

### Regulation of glucose metabolism by miR-31

We next explored in detail how miR-31 expression regulates the above identified metabolic pathways. To analyse the role of miR-31 expression in glucose metabolism (Fig. 2a), we first measured uptake of the fluorescent glucose compound 2-[N-(7-nitrobenz-2-oxa-1,3-diazol-4-yl)amino]-2-deoxyglucose (2-NBDG) as a proxy for glucose uptake (Extended data Fig. 2a) and observed on average a reduction of 2-NBDG uptake of 20%-30% after ectopic miR-31 expression (Fig. 2b) This reduction was observed for several cell lines including 2 additional keratinocyte cell lines (CCD-1106(Hsieh *et al*, 2014) and CCD-1102KERTr). Expressing an anti-miR for miR-31 did not, or only mildly increase 2-NBDG fluorescence (Fig. 2b) possibly reflecting a low level of endogenous miR-31 expression. Furthermore, miR-31- induced reduction of 2-NBDG fluorescence was independent of glucose concentration (Extended data Fig. 2b) and also growth factor-induced 2-NBDG fluorescence was reduced to similar extent (Extended data Fig. 2c). These findings are consistent with miR-31 repressing glucose uptake. In keratinocytes GLUT-1 acts as the major glucose transporter(Zhang *et al*, 2018) and our proteomics data showed reduced expression of GLUT-1 despite lacking an obvious miR-31 seed sequence in the 3’UTR. We confirmed reduced GLUT-1 expression following miR-31 expression by immune fluorescence microscopy (Extended data Fig. 2d). GLUT-1 is a low Km and ATP-independent glucose transporter and hexokinase (HK) mediated glucose-6-phosphate formation is the rate limiting step for glucose utilization(Ishihara *et al*, 1994). We observed miR-31 induced reduction in mRNA expression of HK1 and HK2 (Fig. 2c) and this corroborates the observed reduction in glucose-6-Phosphate (Fig. 1d). In addition, expression of all three Phospho-Fructose Kinase isozymes is reduced upon miR-31 expression (PFKM, PFKL, PFKP (Fig. 2c). Combined with the reduced expression of Aldolase (ALDOC, Fig. 1c) this suggests that miR-31 expression reduced glucose uptake also results in decreased levels of glycolytic intermediates and possibly reduced glycolysis. However, when measuring extracellular acidification rate under normal culture conditions using Seahorse technology, we do not observe a significant miR-31 induced change in lactate production (Fig. 2d). Importantly, under limiting glucose conditions, miR-31 expression significantly enhanced lactate production upon glucose addition (Fig. 2e upper panel). Our SILAC data indicate that miR-31 expression also reduced Pyruvate dehydrogenase complex member X (PDHX) and Pyruvate Carboxylase (PC) expression (Fig. 1c) and this was confirmed by western blotting (Fig. 2f). Both enzymes regulate entry of pyruvate into the mitochondria and therefore decreased expression may divert pyruvate flux towards lactate. In agreement, under low glucose conditions similar to miR-31 expression, pharmacological inhibition of pyruvate entry by UK-5099 a potent inhibitor of the mitochondrial pyruvate carrier (MPC1, Fig. 2g upper panel) as well as siRNA-mediated knockdown of PC increased glycolytic flux towards lactate after glucose addition (Fig. 2h upper panel). This suggests that miR-31 despite reducing glucose uptake maintains glycolytic flux towards lactate by reducing pyruvate entry into mitochondria.

**Figure 2.**
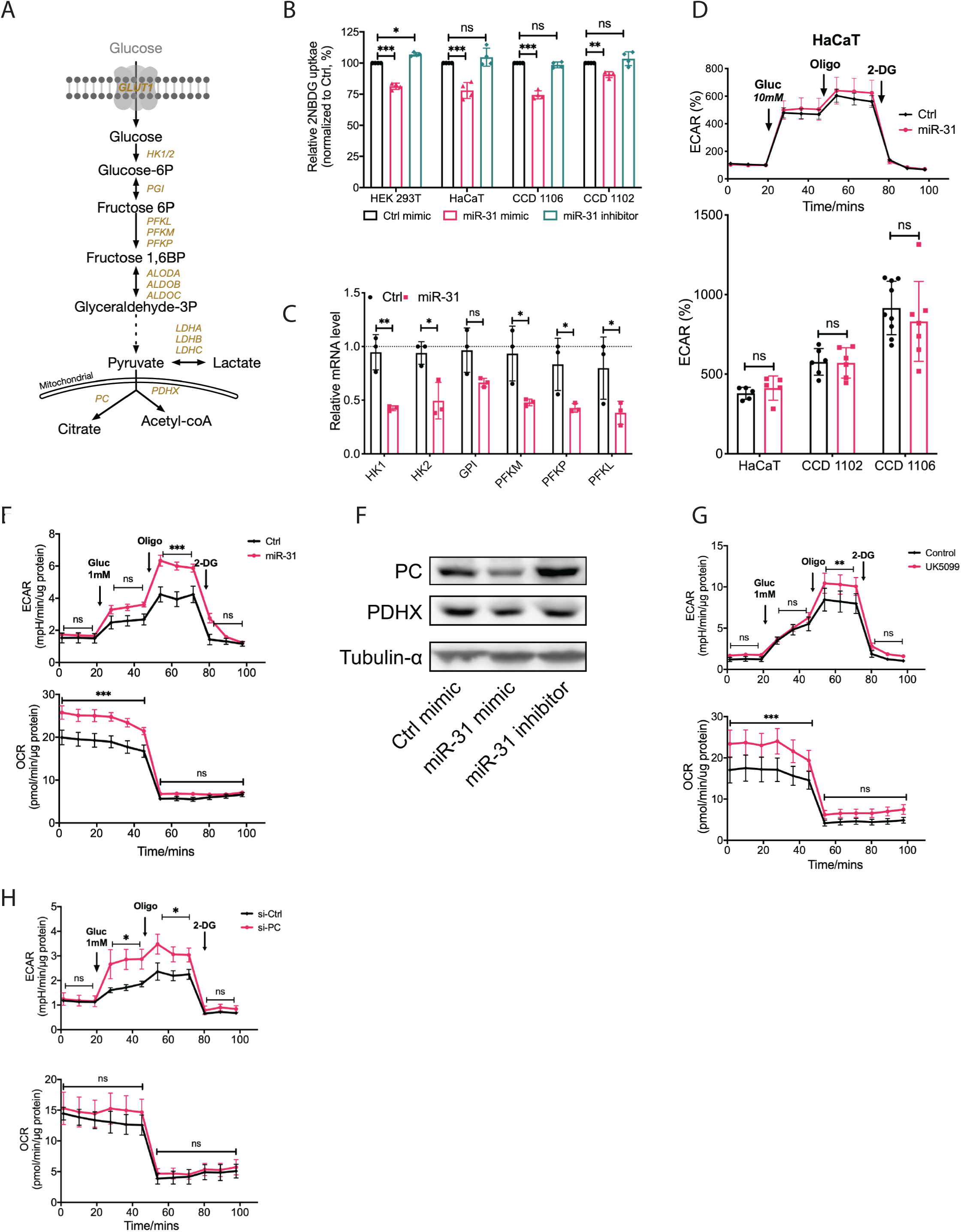
**A. Schematic picture of the glycolysis pathway.** Genes regulating the various metabolite conversions in glycolysis, for which protein expression changes after miR-31 overexpression are indicated. **B. miR-31 affects glucose uptake**. Measurement of 2-NBDG uptake in different cell lines by flow cytometry. n=3 replicates, data are represented as mean± SD, and are compared to ctrl, *, p<0.05; **, p<0.01; ***, p<0.001. **C. miR-31 affects mRNA expression of glycolytic enzymes**. qRT-PCR analysis of glycolytic gene expression in HaCaT cells upon miR-31 overexpression. β-actin was used as a house-keeping control gene, n=3, data are represented as mean± SD, and compared to ctrl, *, p<0.05; **, p<0.01. **D. miR-31 expression does not affect glucose to lactate flux under normal glucose condition.** Standard glycolysis stress test (10 mM glucose) of HaCaT cells treated with miR-31 overexpression (upper panel). Extracellular acidification rate (ECAR) was baselined to third measurement. Gluc, glucose; Oligo, oligomycin; 2-DG, 2-Deoxy-D-glucose. Results from three different cell lines were summarized in bottom panel. n=5-9, data presented as mean± SD. **E. miR-31 expression increases lactate production under low glucose condition**. Low glucose addition (1 mM) of glycolysis stress test of HaCaT cells upon miR-31 overexpression. ECAR (up panel) and oxygen consumption rate (OCR, bottom panel) were normalized to protein concentration. n=5, data presented in mean± SD, and compared to ctrl, *** means p<0.001. **F. mir-31 lowers PC and PDHX expression**. Western blot analysis of PC and PDHX expressions in HaCaT cells, Tubulin α was used as loading control. **G-H. Downregulation of PC and PDHX phenocopies miR-31**. Low glucose addition (1 mM) in glycolysis stress test of HaCaT cells upon UK5099 (an inhibitor of mitochondrial pyruvate carrier, G) or pyruvate carboxylase siRNA (si-PC, H) treatments. ECAR and OCR were normalized to protein concentration. n=4, data presented in mean± SD, and compared to ctrl, *, p<0.05; **, p<0.01; ***, p<0.001. p values (indicated about relevant comparison) were calculated by Two-way ANOVA with Sidak test (B-E, G and H).

Alongside with increasing lactate production following glucose stimulation, miR-31 expression increased basal mitochondrial O_2_ consumption in the absence of glucose, yet this was not further increased by glucose addition. Importantly, this result is identical to UK-5099 mediated MPC inhibition (Fig. 2g lower panel) and siRNA-mediated knockdown of PC showed a similar trend (Fig. 2h lower panel). The latter is in agreement with the study of Cheng et al. showing that PC levels and activity regulate the choice between glucose and glutamine-mediated anaplerosis(Cheng *et al*, 2011), whereby reduced PC increases glutamine-dependent anaplerosis. Taken together, miR-31 expression reduced glycolytic flux towards mitochondrial metabolism, whilst upholding lactate production, and miR-31 induced an increase in mitochondrial O_2_ consumption.These results thus suggest that miR-31 induces mitochondrial activity through alternative sources, like glutamine metabolism.

### Regulation of glutamine metabolism by miR-31

Our proteomic analysis (Fig. 1c) showed a strong downregulation of glutamine synthase (GS) following miR-31 expression making GS a prime target in mediating possible miR-31-induced glutamine-dependent anaplerosis (schematic representation Fig. 3a). Reduced GS expression induced by miR-31 was confirmed by immunoblotting (Fig. 3b). In addition, to GS regulation, we also observed a small increase in expression of glutamate-oxaloacetic transaminase (GOT1) and the mitochondrial aspartate/glutamate transporter (AGC1/SLC25A12) (Fig. 3b). Consequently, miR-31 expression increased glutamine uptake and apparently resulting in a surplus of glutamate production as we observed profound glutamate secretion (Fig. 3c). Glutamine to glutamate conversion requires glutaminase (GLS) and CB-839 is a specific pharmacological inhibitor of GLS. Inhibition of GLS reduced glutamine uptake but did not inhibit miR-31 induced increase, but fully inhibited glutamate secretion in the presence or absence of miR-31 (Fig. 3c). Glutamate can be exported through various transporters including the cystine-glutamate antiporter (xCT), a heterodimer consisting of SLC7A11 and SLC3A2. miR-31 enhances mRNA expression of both components (Extended data Fig. 3a). However, cystine uptake is only slightly increased by miR-31. This may suggest that additional glutamate transporters are being used (e.g., SLC1A3 or SLC1A4) and that usage of the xCT antiporter is determined by conversion of glutamate to glutathione and that the observed miR-31 induced reduction in glutamate-cysteine ligase modifier (GCLM, Fig. 1c) constrains cystine uptake. Next, we tested a role for miR-31 in glutamine-dependent anaplerosis under low glucose condition. As shown, miR-31 expression increased mitochondrial oxygen consumption under low glucose conditions and this was dependent on extracellular glutamine (Fig. 3d). Glutamate cannot replace glutamine in supporting miR-31 dependent mitochondrial metabolism (Extended data Fig. 3b) or miR-31 induced lactate production. Furthermore, the miR-31-induced increase in both mitochondrial metabolism and lactate production was inhibited by CB-839 treatment (Fig. 3e). Substrate supply for OXPHOS is controlled in many ways and the Malate/Aspartate (MAS) shunt plays a role by regulating pyruvate supply and the NAD/NADH balance between cytosol and mitochondria. Increased expression of AGC1/SLC25A12 and GOT1 following miR-31 expression suggests also a key role for MAS in miR-31 induced mitochondrial metabolism and lactate production. Indeed, AGC1/SLC25A12 knockdown reduced miR-31 driven mitochondrial metabolism and lactate production (Fig. 3f). Combined these results indicate that miR-31 stimulates anaplerosis through oxidative glutaminolysis and stimulates aspartate transport to the cytosol to generate pyruvate.

**Figure 3.**
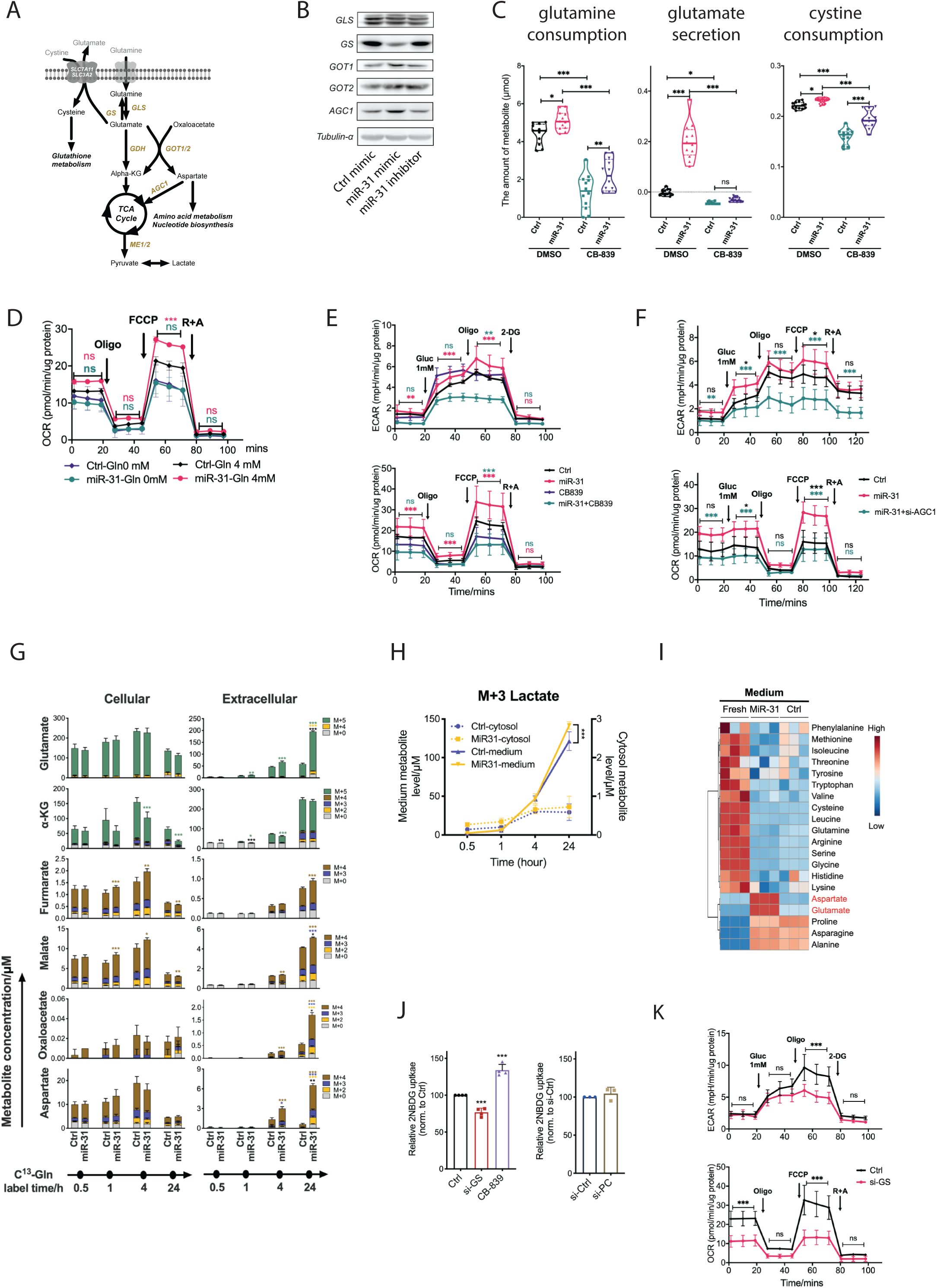
**A. Schematic diagram of glutaminolysis pathway**. TCA cycle, tricarboxylic acid cycle; Alpha-KG, alpha-ketoglutarate; GS, glutamine synthetase; GLS, glutaminase; GDH, glutamate dehydrogenase; GOT1/2, Glutamic-oxaloacetic transaminase 1/2; AGC1, aspartate/glutamate transporter; ME1/2, malic enzyme 1/2. **B. miR-31 regulates protein expression of enzymes involved in glutaminolysis**. Western blot analysis of GLS, GS, GOT1, GOT2, AGC1 in HaCaT cells, Tubulin α was used as loading control. **C. miR-31 regulates glutaminolysis**. Violin plot of glutamine consumption, glutamate secretion, and cystine consumption of HaCaT cells upon different treatments. Metabolites were detected by targeted LC-MS metabolomics and normalized to fresh medium for the calculation of consumption or secretion. n=12, *, p<0.05; **, p<0.01; ***, p<0.001. **D. miR-31 regulates mitochondrial metabolism through glutaminolysis**. Mitochondrial stress test of HaCaT cells with miR-31 treatment. Cells were pre-cultured in medium with or without glutamine overnight. OCR normalized to protein concentration. FCCP, Carbonyl cyanide 4-(trifluoromethoxy)phenylhydrazone; R+A, antimycin and rotenone. n=3, statistic result marked with green means difference between miR-31-Gln 0 mM and Ctrl-Gln 0 mM groups; marked with pink represent comparison of miR-31-Gln 4 mM vs Ctrl-Gln 4 mM; **, p<0.01; ***, p<0.001. **E-F. Requirement for GLS and AGC1 for miR-31 regulated glutamine-dependent mitochondrial metabolism.** Glycolysis (upper panel) and mitochondrial (bottom panel) stress tests of HaCaT cells upon CB839 (an inhibitor of GLS, E) and AGC1 siRNA (si-AGC1) treatments under low glucose (1mM) supplementation. n=3-5, statistic result marked with pink represents comparison of miR-31 and Ctrl (E); green(E), CB839 vs miR-31+ CB839; green (F), miR-31 vs miR-31+ si-AGC1; black (F), miR-31 vs black; *, p<0.05; ***, p<0.001. **G. miR-31 directs glutamine flux to aspartate**. Cellular and extracellular levels of metabolites were detected by targeted metabolomics using C13 glutamine tracing protocol in a time course. n=3, data presented in mean± SD, compared to ctrl, *, p<0.05; **, p<0.01; ***, p<0.001. **H. miR-31 enhances lactate labeling derived from glutamine.** The dashed and solid lines represent the intracellular and extracellular lactate levels under different intervention conditions, respectively. n=3, data presented in mean± SD, compared to ctrl-medium, ***, p<0.001. **I. miR-31 increases extracellular glutamate and aspartate**. Heatmap representation of relative amino acid concentrations in cell culture medium after miR-31 overexpression. **J. Glutamine uptake regulates glucose uptake**. 2-NBDG intensity was measured by flow cytometry for the comparison of glucose uptake in HaCaT cells with different treatments. n=3, data represent mean± SD, and compared to ctrl, ***, p<0.001. **K.** GS knockdown is not sufficient to phenocopy metabolic regulation of miR-31. Glycolysis (upper panel) and mitochondrial (bottom panel) stress tests of HaCaT cells upon GS siRNA (si-GS) treatments under low glucose (1mM) supplementation. n=5, data are represented as mean± SD, compared to si-ctrl, ***, p<0.001. p values (indicated for relevant comparison) were calculated by One-way ANOVA with Sidak test (C and I), Two-way ANOVA with Sidak test (D-G), or Student’s t-test (I).

To further corroborate our observations, we performed 15N,13C-glutamine tracing (Fig. 3g). As we predominantly observe miR-31 to stimulate glutaminolysis under low glucose conditions (Fig. 3d) we performed tracing under glucose-free conditions and added low amount of unlabelled glutamine to prevent cell death before adding tracer glutamine. This resulted in rapid (within 30 minutes) labelling of most metabolites. In agreement with lactate production being maintained by miR-31 (Fig 2d) we observed that miR-31 increased lactate labeling derived from glutamine (Fig 3h), indicating that lactate production is the consequence of a combination of increased glutaminolysis and reduced pyruvate entry into mitochondria. We observed additional miR-31 induced increase in cellular malate and fumarate labelling showing miR-31 induced anaplerosis of the TCA cycle. More strikingly, we observed a strong increase in miR-31 induced secretion of not only glutamate but also aspartate, oxaloacetate and to lesser extend malate and fumarate. Glutamate and aspartate secretion following miR-31 expression was independently confirmed using a LC-MS platform for analysis of biogenic amines (Fig. 3i). These results combined suggest that miR-31 expression results in increased glutamine to glutamate conversion by inhibiting GS, however the fate of glutamate is determined by miR-31 dependent regulation of other metabolic enzymes like PC/PDHX and GCLM/AGC1. Together this results in miR-31 facilitated glutamine-dependent anaplerosis under glucose limiting conditions and this produces aspartate, but also that a significant part of the metabolites (aspartate, glutamate but also oxalo-acetate) is secreted (schematic representation Extended data Fig. 3c).

To determine whether indeed next to GS regulation, the regulation of these other miR-31 targets is required to determine its metabolic phenotype we addressed whether reduced GS expression by itself is sufficient to mimic miR-31 expression in regulating glutaminolysis. Interestingly, siRNA-mediated reduction in GS expression, similar to miR-31 expression, also decreased glucose uptake (Fig. 3j), whereas reducing expression of PC did not affect glucose uptake. In contrast to miR-31 expression, siRNA mediated knockdown of GS resulted in decreased glycolysis towards lactate and also decreased mitochondrial metabolism (Fig. 3k). As individual regulation of these miR-31 targets does not phenocopy miR-31 induced metabolism, these results strongly imply that the mechanism whereby miR-31 couples glucose and glutamine metabolism results from combined miR-31-dependent regulation of GS with PC/PDHX and GCLM/AGC1.

### Consequence of metabolic regulation on proliferation and inflammation

Psoriasis and comparable skin diseases emerge through a combination of hyperproliferative keratinocytes and local inflammation. To test consequences of metabolic rewiring by miR-31 expression on disease parameters we first tested cell growth of HaCAT cells under limiting glucose or glutamine conditions. Under normal culture conditions miR-31 expression did not affect cell proliferation, but miR-31 expression reduced cell proliferation under glutamine restricted conditions and in contrast enhanced cell proliferation under glucose limiting conditions (Fig. 4a). Mechanistically, these effects on cell proliferation can be explained by miR-31 induced reduction of GS and consequent enhanced glutaminolysis, but also by reduced PC expression following miR-31 expression, which renders cells more dependent on glutamine for anaplerosis(Cheng *et al*., 2011). In contrast, siRNA-mediated reduction of GLS expression or pharmacological inhibition by CB-839 makes cells dependent on glucose for anaplerosis(Cheng *et al*., 2011). In agreement, we observed synthetic lethality in HaCAT cells between miR-31 expression and CB-839 treatment, whereas anti-miR-31 expression, increased survival after CB-839 treatment (Fig. 4b and Extended data 4a). This GLS dependence of miR-31 expressing cells further corroborates that upon miR-31 expression cells move from endogenous glutamine production to exogenous glutamine uptake as a source of glutamine.

**Figure 4.**
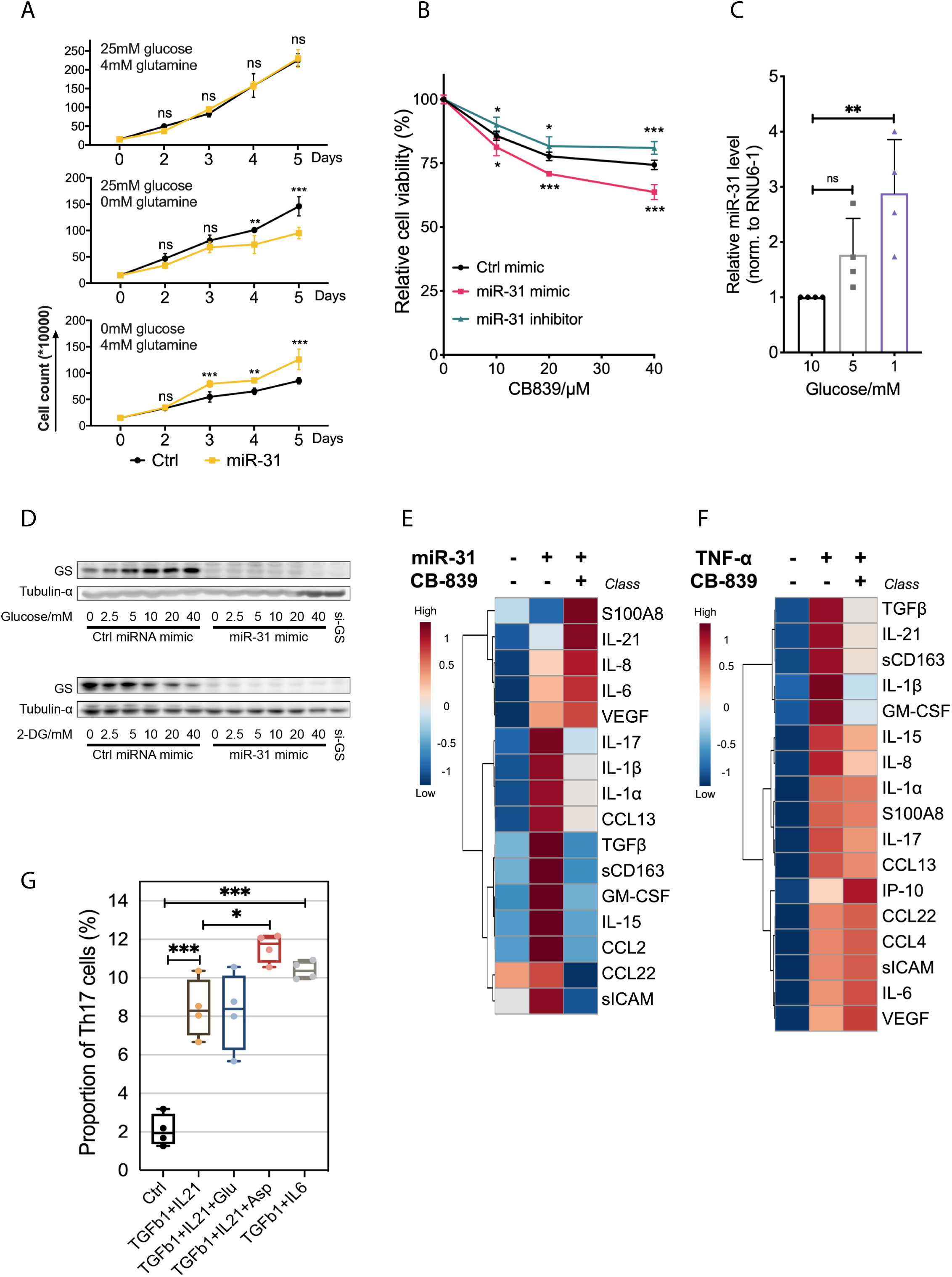
**A. miR-31 regulates cell growth under limiting glucose condition**. Cell growth curves of HaCaT cells with or without miR-31 overexpression in medium with or without glutamine or glucose as indicated. n=5, data showed in mean ± SD, compared to ctrl, **, p<0.01; ***, p<0.001. **B. GLS inhibition impairs viability of miR-31 expressing cells**. Cell viability of HaCaT cell was measured by MTS assay. n=4, data showed in mean ± SD, compared to ctrl, *, p<0.05; **, p<0.01; ***, p<0.001. **C. miR-31 expression is regulated by glucose levels**. qRT-PCR analysis of miR-31 expression in HaCaT cells cultured at different glucose concentrations in the culture medium. RNU6-1 was used as house control gene. n=4, data are represented as mean ± SD, compared to ctrl, **, p<0.01. **D. The miR-31 target GS is regulated by glucose**. GS expression in HaCat cells was determined by immunoblotting of total cell protein lysates. Cells expressed control mimic miR or miR-31 and were cultured in medium with indicated glucose concentration (upper panel) or glycolysis was blocked using different concentrations of 2-deoxy-glucose as indicated. Tubulin α was used as control for equal loading of protein. **E-F. miR-31-dependent glutamine metabolism regulates expression of a subset of cytokines.** Heatmap representation of cytokine expression in HaCaT cells induced upon miR-31 expression (E) or TNFa treatment (F). CB-839 treatment was used to determine dependency on extracellular glutamine usage for expression of cytokines. **G. IL-21, TGFb1, and aspartate enhance Th17 cell differentiation.** The proportions of CD4+CD25-RORγt+Th17 cells in CD4+CD25- T cells were analysed by FACS. Asp, aspartate; Glu, glutamate. p values (indicated about relevant comparison) were calculated by Two-way ANOVA with Sidak test (**A** and **B**) or One-way ANOVA with Sidak test (**C** and **G**).

As the miR-31 effect on cell proliferation is dependent on the consequent metabolic changes we addressed whether metabolic changes in reverse affect miR-31 expression and interestingly, we observed that glucose restriction by itself induces expression of miR-31 (Fig. 4c) and also reduced GS expression (Fig. 4d). Together this suggests that glucose availability can couple in an inverse manner to glutaminolysis through regulating expression of miR-31 and consequent expression of GS and other miR-31 targets.

Crosstalk between keratinocytes and immune cells mediated by cytokines is important in maintaining the diseased state (reviewed in (Lowes *et al*, 2014)). Previous studies(Xu *et al*., 2013; Yan *et al*, 2015) identified miR-31 as a pro-inflammatory factor in psoriasis, showing that miR-31 activated the NF-kB pathway and thereby increased secretion of a set of cytokines. Therefore, we next determined a possible role for miR-31 induced metabolism in the expression of mediators of the inflammatory response in HaCAT cells. Expression of miR-31 by itself induced expression of various cytokines most notably in the context of psoriasis IL-17 and IL1b (Fig. 4e, Extended table 4). Anti-IL17 treatment is currently used for psoriasis treatment and interestingly expression of IL-17, IL1b, and other, but not all cytokines were inhibited by CB-839 treatment. Thus, glutamine metabolism mediates in part miR-31 induced regulation of these cytokines. Interestingly CB-839 treatment also enhanced expression of some of the miR-31 induced cytokines and this may indicate that these are sensitive to glycolytic metabolism. In psoriasis TNFa is an important inflammatory cytokine and NF-kB activator and in HaCAT cells a subset of TNFa- induced cytokines overlaps with those regulated by miR-31, most importantly in the context of psoriasis TGFb and IL1b (Fig. 4f, Extended table 5). Most notably, TNFa-induced expression of these and some others is sensitive to CB-839 treatment indicating the relevance of miR-31 controlled glutamine metabolism in the inflammatory control of psoriasis.

### MiR-31 induced secretome induces Th17 differentiation

The above defines a miR-31 induced secretome that combines specific metabolites as well as cytokines and partially depends on the regulation of glutamine metabolism by miR-31. To test a functional consequence of the miR-31 induced secretome we tested differentiation of naive T-cells towards Th17 cells, as this T-cell subtype specifically marks psoriasis.

Naive T-cells were treated with different cytokines and we observed that combined with TGFb1 both IL-6 and IL-21 induce Th17 differentiation and importantly this can be further increased by adding aspartate to the medium whereas glutamate did not induce additional differentiation. Recently, it has been shown that glutaminolysis in mice stimulates Th17 differentiation (Johnson *et al*, 2018; Kono *et al*, 2018) and therefore the latter result was unexpected. However, high extracellular glutamate, inhibits cystine uptake by the cystine/glutamate antiporter (system xCT (Dixon *et al*, 2012) and consequently results in increased levels of ROS, that even impair Th17 formation(Johnson *et al*., 2018). In this respect glutamate may not simply replace glutamine.

Taken together these results indicate that miR-31 expression in keratinocytes also supports in part the inflammatory response by stimulating Th17 differentiation.

### Role for miR-31-induced metabolic regulation in psoriasis

Nutrient availability is likely not homogenous in the skin, with only the basal layer being adjacent to blood vessels and upper layers therefore relatively devoid of nutrients. This would be exaggerated during skin diseases characterized by keratinocyte hyperproliferation. We used immunohistochemistry to study whether the miR-31-mediated switch to glutaminolysis and its effect on cell proliferation under glucose limiting conditions in vitro is relevant to psoriasis.

The biopsies of psoriasis patients used for IHC show typical epidermal thickening (Fig. 5a) increased proliferation (Ki67 staining, Fig. 5b) and displayed increased miR-31 expression specifically in the spinous layer (Fig. 5c). Next, we tested expression GLUT-1, GLS and GS (Fig. 5d). In normal skin GLUT-1 expression appears mostly restricted to cells present in the basal layer and expression is lost when cells are moving upwards into the spinous and granular layer. Also, in psoriatic lesions from human patients GLUT-1 expression appears mostly restricted to cells in the basal layer and strongly reduced in the upper layers (Fig. 5d for quantification see Extended data Fig. 5a). In contrast to GLUT-1, GLS staining is high in all cell layers both in normal skin as well as in psoriatic skin. GS expression was low in normal skin but increased in cells of the basal layer of the skin in psoriatic lesions. However, in psoriatic lesions GS expression was decreased in the spinous and granular layer concordant with miR-31 regulating GS and expression of miR-31 being low in the basal layer as compared to an increase in the spinous and granular layer. Taken together these data suggest that the skin displays metabolic compartments whereby the basal layer is GLUT-1 positive and the upper layers are GLUT-1 negative, but GLS positive (Fig. 5e).

**Figure 5.**
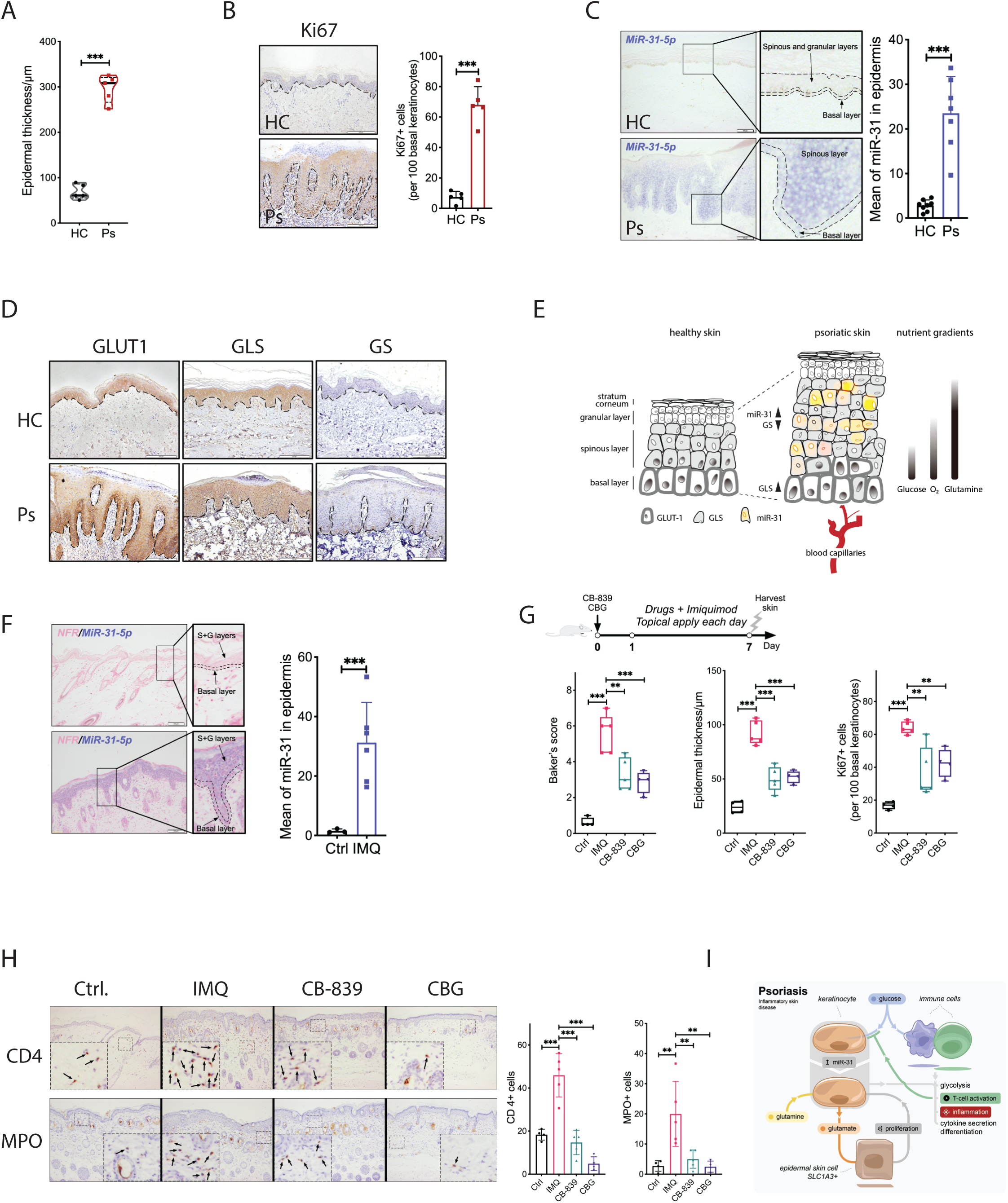
A-D. **Characterization of psoriasis parameters in patient cohort.** A. Violin plot of epidermal thickness of skin biopsies taken from healthy people (HC) and psoriasis patients (Ps). n=5, ***, p<0.001. B. Cell proliferation as assessed by Ki67 immunostaining of skin from HC and Ps. Number of Ki67 positive staining cell was counted as for quantification (right panel). Dashed lines indicate the boarder of epidermis and dermis respectively. n=5, data are represented as mean ± SD, compared to ctrl, ***, p<0.001. C. Representative pictures of in situ hybridization (ISH) staining of miR-31 in skin biopsies from HC (n=8) and Ps (n=7). Quantification of results showing the average level of miR-31 in skin epidermis (right panel), mean ± SD , ***, p<0.001. D. GLUT1, GLS, and GS staining of human skin revealed a different metabolic signature between Ps and HC. Dashed line indicates the boarder of epidermis and dermis. n=5 in each group. E. **Schematic picture illustrating expression distribution in the epidermis of miR-31, GLUT1, GLS and GS of normal skin versus psoriatic skin.** F. **Distribution of miR-31 expression in IMQ model skin**. Representations of in situ hybridization staining (ISH) of miR-31 in skin biopsies from mouse of normal (Ctrl, n=3) and imiquimod-induced psoriatic mouse model (IMQ, n=6). Quantification result showing the average level of miR-31 in epidermis(right panel), mean ± SD , ***, p<0.001. **G-H. GLS inhibition alleviates psoriasis pathology.** G. Disease activity was evaluated by Baker’s score, epidermal thickness, and Ki67 positive cells in basal layer of epidemics. n=4-5, mean ± SD , **, p<0.01; ***, p<0.001. H. **GLS inhibition reduces the number of immune cells**. Representative pictures of Immunohistochemistry staining of CD4 (upper panel) and MPO (bottom panel) in skin biopsies from mouse (n=4-5). Quantification result showing the average number of CD4 positive or MPO positive cells in skin (right panel), mean ± SD , **, p<0.01 , ***, p<0.001. I. **Model summary of cell coexist through metabolism reprogramming in different disease.** p values (indicated about relevant comparison) were calculated by Student’s t-test (A-C, F), and One-way ANOVA with Sidak test (G and H).

To address causality, we made use of the IMQ mouse model for psoriasis. The IMQ model employed, similar to human patients, showed increased scaling (Extended data Fig. 5b) and a gradual increase in epidermal thickness (Extended data Fig. 5c), but also an increase in miR-31 expression in the spinous layer (Fig. 5f). Also, the pattern of GLS and GLUT-1 expression in this model reflected human disease (Extended data Fig. 5d). As we observed in cell culture synthetic lethality between miR-31 expression and GLS inhibition by CB-839, we tested the effect of CB-839 treatment in this model for psoriasis. Cutaneous application of CB-839 was compared to treatment with Calcipotriol and Betamethasone Dipropionate Gel (CBG, commercial name Xamiol), a commonly used treatment option for psoriasis(Kuehl & Shear, 2018). CB-839 treatment decreased epidermal thickness, decreased keratinocyte proliferation as measured by Ki67 staining and lowered the histopathological score based on baker’s system (Fig. 5g). Furthermore, we observed a significant reduction in the number of capillary vessels (Extended data Fig. 5e), following CB-839 treatment. In line with a role for glutamine metabolism regulating cytokine production by keratinocytes (Fig. 4e-4f) we also observed a reduction in CD4+ T-cells and Myeloperoxidase positive (MPO+ cells; neutrophil granulocytes) indicating reduced inflammatory response (Fig. 5h).

Taken together CB-839 showed similar or better therapeutic effect compared to CBG/Xamiol in alleviating psoriatic symptoms in the IMQ model. Importantly, IHC analysis showed that both CB-839 and CBG/Xamiol treatment in the IMQ model also partially normalized the expression pattern of GLUT-1, GLS and GS (Extended data Fig. 5f).

## Discussion

Psoriasis is one of the most common human skin diseases characterized by excessive growth and aberrant differentiation of keratinocytes combined with a strong local inflammatory immune response. Many studies have addressed the genetic underpinning of psoriasis including the expression and potential role of aberrant miRNA expression. Here we confirm high miR-31 expression in human biopsies of psoriasis and in the IMQ mouse model system for psoriasis. We present evidence that miR-31 expression induces metabolic rewiring through a combined deregulation of enzymes involved in glycolysis, glutaminolysis and lipid metabolism. We observed that miR-31 expression limits glucose flux towards mitochondrial metabolism whereas it upholds lactate production and increases mitochondrial respiration. Lactate production is maintained in part by reducing pyruvate entry into mitochondria through reduced PC and PDHX expression and in part through increased glutaminolysis, which also maintains mitochondrial respiration. miR-31 regulation of glutaminolysis occurs at multiple levels. Primarily, expression of miR-31 reduced GS expression resulting in increased glutamine uptake and excess glutamate. Subsequently, miR-31 expression steers glutamate metabolism through the combined regulation of several metabolic enzymes/pathways. First, miR-31 expression increased glutamate secretion by regulation of the xCT antiporter (increased expression of SLC7A11 and SLC3A2), likely in combination with downregulation of GCLM. Further, glutamate metabolism is determined by miR-31 dependent regulation of the mitochondrial Malate-Asparte shuttle (MAS). Interestingly we observed regulation of GOT1, MDH1, ME1 and SLC25A12 as compared to GOT2, MDH2, ME2 and SLC25A11, indicating that glutamine-directed anaplerosis of the TCA cycle additionally results in MAS-dependent secretion of aspartate and oxaloacetate into the cytosol and this is corroborated with the observed increase in these metabolites when measuring flux undr glucos limiting conditions (see for schematic overview Extended data Fig. 3c). Importantly, the mode of miR-31 dependent regulation of glutaminolysis allows cell survival under glucose limiting conditions.

Inhibition of glutaminolysis by CB-839 treatment mitigates most of the effects of miR-31 on glutamine metabolism suggesting that miR-31 redirects cells from endogenous glutamine production to exogenous glutamine uptake as a source of glutamine.

In keeping, with our results showing differential expression of miR-31 and GLS/GS within the layers of healthy and psoriatic skin, we find that CB-839 treatment is effective in alleviating psoriatic disease in the IMQ mouse model.

Many studies have reported a possible correlation between metabolic syndrome, diabetes and the incidence of skin disease such as psoriasis. However, a recent meta-analysis covering 26 clinical studies concluded that there is no clear support to suggest a link between defective glucose metabolism and psoriasis(Friis *et al*, 2019). In contrast others(Zhang *et al*., 2018) and our results clearly provide support for the involvement of dysregulated metabolism in psoriasis, but not as a predisposition but rather as part of the mechanism that drives the disease when manifest.

Our results identify inhibition of glutaminolysis as a treatment opportunity for psoriasis. Currently, various psoriasis treatment regimens are available. Most recently, these are drugs targeting the IL-23/IL-17A pathway which are reasonably effective in relieving skin symptoms of psoriasis. However, more general treatment options such as fumarate and methotrexate are still clinically used(Mrowietz *et al*, 2018; Wang & Tsai, 2017). The mode of action of fumarate and its related compound dimethyl-fumarate in relieving psoriatic symptoms remains unclear (Mrowietz *et al*., 2018). Interestingly, recently GAPDH has been suggested to be targeted by fumarate (Kornberg *et al*, 2018). In addition, GLUT-1 targeting has been proposed as treatment option for psoriasis (Zhang *et al*., 2018). Methotrexate targeting Di-Hydro-Folate reductase (DHFR) inhibits nucleotide synthesis and thus its efficacy indicates limitations with respect to nucleotide synthesis in psoriasis. Aspartate is required for pyrimidine nucleotide metabolism and in addition has been shown to be rate limiting for proliferation under hypoxia. This corroborates our proposed role for miR-31 upregulation and regulation of aspartate metabolism in driving keratinocyte proliferation in psoriatic disease.

In cancer, stem cells are considered the cell of origin and a recent study showed that the glutamate/aspartate transporter SCL1A3/EAAT3 marks epidermal stem cells(Reichenbach *et al*, 2018). In psoriasis, to our knowledge, a cell of origin has not been defined but, as psoriasis similar to cancer entails a disease of aberrant cell proliferation it is well conceivable that in psoriasis hyperproliferation may originate from or driven by SCL1A3 positive stem cells. This concept would be of interest as it further emphasizes the potential importance of miR-31 induced glutamate/aspartate metabolism, in the sense that miR-31 expressing keratinocytes not only affect immune cell function but may also drive stem cell function. Clearly this requires further study.

In humans, TH17 cell differentiation is induced by IL-1b as well as IL-23, and possibly transforming growth factor-beta (TGFb) in the presence of inflammatory cytokines such as IL-6, IL-21 and IL-23 (reviewed in (van den Berg & McInnes, 2013)). Interestingly, we show that production of TGFb and IL-1b by keratinocytes depends on miR-31 expression indicating a potential role for miR-31 in instructing TH17 differentiation. In addition, recent years have shown the importance of metabolism in defining T-cell identity and function (reviewed in e.g., (Buck *et al*, 2017)). In general, increased glycolysis is essential for T-cell activation and consequently proliferating cells and immune cells compete with respect to glucose availability. In cancer it has been suggested that this metabolic competition is to the advantage of the proliferating cancer cells displaying a Warburg phenotype, whereby immune cells are under metabolic constraint owing to the lack of glucose to sustain their metabolism (Chang *et al*, 2015). This constraint contributes to the lack of an endogenous immune response towards cancer. In contrast, skin disease, including psoriasis, is characterized by proliferation of keratinocytes and immune cell activation. By analogy to the aforementioned the proliferating keratinocytes are likely metabolically constrained and this maybe be exaggerated by the skin environment. GLUT-1 expression is only evident in the basal layer suggesting that glucose availability is restricted in upper dermal layers. Interestingly, we observed that decreasing glucose levels resulted in increased miR-31 expression and this is corroborated by our observation that miR-31 expression in the IMQ model is observed also in the upper dermal layers. Together this suggests that the metabolic constrain imposed by immune cell activation and the skin environment actually induces increased miR-31 expression and thereby propels keratinocyte proliferation. How glucose availability intersects with other proposed regulatory pathways of miR-31 expression (Xu *et al*., 2013; Yan *et al*., 2015) remains to be investigated. Nutrients and O2 are provided by vasculature present below the basal layer and as such a gradient of nutrients upwards will emerge. Similar to glucose there will be limited O2 available and skin hypoxia along the same line of reasoning may be an important consequence of psoriasis. Hypoxia inhibits the mitochondrial electron transport chain and thereby aspartate synthesis(Garcia-Bermudez *et al*., 2018). Consequently, aspartate levels will decrease under low oxygen. As aspartate is an essential metabolite in nucleotide synthesis, cell proliferation can therefore only occur when aspartate levels are increased either through miR-31 expression as shown here or through import by e.g., SLC1A3.

anti-IL17 therapy is highly effective in psoriasis patients indicating the importance of IL-17 dependent immune cells like Th17 and γδT17 in psoriasis. Glutaminolysis has been reported to control Th17 differentiation in mice (ref) and in agreement herewith it has been shown that also the Th17 cells in psoriasis display increased glutaminolysis, likely through increased GLS1 expression in these cells (Johnson et al., 2018; Kono et al., 2018). Here we show that the involvement of glutamine metabolism in psoriasis acts beyond the differentiation of Th17 cells and is also part of the mechanism whereby keratinocytes contribute to this disease.

Based on the requirement for glutaminolysis in Th17 differentiation, Xia et al. (Xia *et al*, 2020) also described the potential of GLS inhibition in the treatment of psoriasis. However, there it was suggested that the effect on psoriasis was due to GLS inhibition in T cells and consequent reduced Th17 differentiation. Our results add to this suggestion and provide novel understanding of inflammatory skin disease progression (summarized in model Fig. 5i). This model illustrates a mode of metabolic co-existence in psoriasis, as opposed to metabolic competition in cancer. For the diseased state this implies that either active immune cells and proliferating cells can coevolve (psoriasis) due to differential metabolic usage or that proliferating cells outcompete activation of immune cells by metabolic competition as in the case of cancer. Finally, this model illustrates how in general cellular heterogeneity in disease combines with the metabolic environment to sustain or enable disease, but also shows that disconnecting this metabolic interaction between cell populations can be effective in treating disease as shown here for psoriasis.

## Acknowledgements

We wish to thank Prof Matthew vander Heiden (MIT, Boston), Dr. Emmerik Leijten (Rheumatology, UMCU) Dr. Prof Nanda Verhoeven-Duif, Dr Judith Jans (Metabolic Genetics, UMCU) Maria Rodriguez-Colman, and other members of the Burgering lab for comments and discussion. Research in the Burgering lab is financially supported by a the Oncode Institute. Research at the Second Affiliated Hospital, Guangzhou University of Chinese Medicine is supported by a grant of the National Natural Science Foundation of China (No.81873302) and 3 grants of Guangdong Provincial Hospital of Chinese Medicine (No. 2019KT311, No. 2019KT312, and No. 2019KT313). Animal experiments were approved by Animal ethics committee of Guangdong Provincial Hospital of Chinese Medicine (China, No. 2018068).

## Materials and methods

### Cell culture

HEK293T and HaCaT cells were cultured in DMEM (Gibco) supplemented with 10% FBS and 1% penicillin–streptomycin. CCD 1102 KERTr (ATCC® CRL-2310™) and CCD 1106 KERTr (ATCC® CRL-2309™) were cultured in Keratinocyte-Serum Free Medium (Gibco 17005-042) with added Keratinocytes Supplements (Gibco 37000-015) including Bovine Pituitary Extract (BPE; Gibco 13028-014) and human recombinant epidermal growth factor (EGF; Gibco 10450-013). All cells were cultured at 37 °C and 5% CO2.

### Glucose uptake measurement

A nonradioactive assay was performed for the measurement of glucose uptake in different cell lines by using 2-NBDG (2-(N-(7-nitrobenz-2-oxa-1, 3-diazol-4-yl) amino)-2-deoxyglucose, Cayman chemicals, USA), a fluorescent deoxyglucose analog. Briefly, cells were incubated with 50μM of 2-NBDG in a cell incubator with 37℃ for one hour. Then, cells were collected and washed by 2 times of PBS. In the end, cells were stained with PI about 5 mins for identification of live cells, and then loaded into Celesta flow cytometer (BD Biosciences) for FACS analysis.

### Proliferation Measurement

For cultured cell proliferation, equal HaCaT cells numbers (150,000 cells per well) were seeded in 6-well plate, overexpressed with miR-31 mimic or scramble miRNA mimic (Ctrl) in indicated medium (complete growth medium with 25 mM glucose and 4 mM glutamine, glutamine starvation medium with 25 mM glucose and without glutamine, and glucose starvation medium with 4 mM glutamine and without glucose, refreshed medium at day 3), harvested at indicated times, and then counted in the presence of Trypan blue (Biorad, 145-0013). For colony-forming assay, HaCaT cells were seeded in 6-well plates and treated with overexpression of miR-31 mimic or inhibitor for 36h, followed with 10 μM CB-839 addition. Cell culture medium and CB- 839 were refreshed every 3day in a total of 14 days treatment. For staining, cells were washed with PBS twice, fixed with 100% methanol 20 mins, and stained with 0.25% Crystal Violet (Sigma, C0775) in 6.25% ethanol for 30min at room temperature, washed 3 times with ddH2O, dried at room temperature, and photographed.

### MTS assay

Cells were plated in 96-well microtiter plate and received different treatments before MTS assay. For the MTS assay, 20ul of MTS reagent was directly added into the culture medium and incubated at 37℃ incubator for 3h. Finally, the formazan dye is quantified by measuring the absorbance at 490nm.

### RNA extraction and quantitative Realtime-PCR

Total mRNA was extracted following the manufacture of Qiagen RNeasy Kit (Qiagen), and reversed transcribed using the iScript cDNA Synthesis Kit (Bio-Rad). qReal-time PCR was performed using SYBR Green FastStart Master Mix (Roche) in the CFX Connect Real-time PCR detection system (Bio-Rad). Data was normalized to beta-actin and presented as a relative expression level that was calculated by using delta delta Ct analysis. All primer sequences of targeted genes, where derived from primerbank (https://pga.mgh.harvard.edu/primerbank/) and described in Extended table 6. For quantification of microRNA expression, mature miR-31 was quantified using MystiCq® microRNA qPCR Assay (Sigma, MIRRM00-100RXN) according to the manufacturer’s instructions. Primer of miR-31 was ordered from Sigma (MIRAP00090) and RNU6-1 was used as the internal control (Sigma, MIRCP00001).

### Stable isotope labelling of amino acids in cell culture (SILAC) experiment setup

For SILAC labelling HaCaT cells were cultured in SILAC-labelled DMEM (Thermo Fisher Scientific) with 10% dialyzed FBS and 1% of penicillin/streptomycin containing L-arginine and L-lysine (light medium) or 13C6-L-arginine and 3C6-L-lysine (heavy medium) for at least 14 days to eliminate non labelled arginine and lysine. The day before the transfection, 1.5x10^5^ cells were seeded in a 6-well plate in a final volume of 3mL. On the next day, the medium was changed and the 60-70% confluent cells were transfected with a specific miRNA mimic either for miR-31 mimic or control miRNA mimic at a final concentration of 30nM together with lipofectamine RNAiMAX and Opti-MEM. 48h post transfection cells were harvested by adding lysis buffer containing 8M Urea, 1M ammonium bicarbonate, 10nM tris (2-carboxyethyl) phosphine, 40nM chloroacetamide. Cell lysates were incubated at 95°C for 5 min, sonicated and diluted to 2M Urea with 1M ABC. After protein quantification using the BCA protein assay, cell lysates from miR-130a or miR-708 transfected cells generated from heavy medium and SCR transfected cells from light medium were mixed 1:1 (reverse mode) and vice versa (forward mode). Proteins were digested overnight with 2% (w/w) trypsin, next peptides were fractionated based on their molecular mass using ultra performance liquid chromatography (UltiMate-3000 system, Thermo Fisher Scientific) and finally desalted and acidified on a C-18 cartridge (3M, Saint Paul, Minnesota, USA). C18-stagetips were activated with methanol, washed with buffer containing 0.5% formic acid in 80% ACN (buffer B) and then with 0.5% formic acid (buffer A). After loading of the digested sample, stage-tips were washed with buffer A and peptides were eluted with buffer B, dried in a SpeedVac, and dissolved in buffer A. Peptides were electro-sprayed directly into an Orbitrap Fusion Tribrid Mass Spectrometer (Thermo Fisher Scientific) and analysed in Top Speed data-dependent mode. Raw files were analysed using Maxquant software. For identification, the human Uniport 2017 was searched with both the peptide as well as the protein false discovery rate set to 1%. Proteins identified were filtered for reverse and decoy hits, standard contaminants and selected to have more than 1 unique or razor peptide by using the Perseus software 1.5.1.6. Heavy/light normalized ratios (ratio reverse and ratio forward) were used to quantify protein expression and were further processed for comparative analysis of differential expression among the conditions.

### Western blot analysis and immunofluorescence

Cells were lysed in sample buffer (0.2% m/v SDS, 10% v/v glycerol, 0.2% v/v β-mercaptoethanol, 60 mmol/L Tris pH 6.8) for protein extraction. Proteins were detected using 7.5%–12.5% SDS-PAGE gels and subsequent Western blot analysis with primary antibodies detected by HRP-conjugated secondary antibody. Unless stated otherwise, alpha-tubulin were used as a loading control. For immunofluorescence, cells were fixed with 3.7% paraformaldehyde in PBS, permeabilized with 0.1% v/v Triton in PBS, and blocked with 2% m/v BSA (Sigma). Cells were subsequently incubated with primary antibody and visualized with Alexa secondary antibody (Sigma).

### Bioenergetics

Seahorse Bioscience XFe24 Analyzer was used to measure extracellular acidification rates (ECAR) in mpH/minute and oxygen consumption rates (OCR) in pmol O2/minute. Cells were seeded in 6-well plates and transfected with indicated miRNA mimic (scramble or miR-31-5p) or indicated si-RNAs (si-PC, si-GS, and si-AGC1) for 36h, or treated with pharmaceutical drugs (CB-839 (10 mM final concentration) or UK 5099 (25 mM final concentration) for 24h. Thereafter, cells were seeded in XF24 polystyrene cell culture microplates (Seahorse Bioscience) at a density of 10.000 cells per well.1 hour before the measurements, culture medium was replaced and the plate was incubated for 60 minutes at 37 ℃. For the mitochondrial stress test, culture medium was replaced for Seahorse XF Base medium (Seahorse Bioscience), supplemented with 20 mM glucose (Sigma-Aldrich), 2 mM L-glutamine (Sigma-Aldrich), 5mM pyruvate (Sigma-Aldrich) and 0.56 μl NaOH (1M). During the test 5 μM oligomycin, 2 μM FCCP and 1 μM of Rotenone and Antimycin A (all Sigma-Aldrich) were injected to each well after 18, 45 and 63 minutes respectively. For the glycolysis stress test, culture medium was replaced for Seahorse XF Base medium, supplemented with 2mM L-glutamine and 0.52 μl/ml NaOH (1M). Sensor cartridges (pre-hydrated in XF calibrant solution overnight in a CO2-free incubator) were loaded with glucose (Port A), oligomycin (Port B), and 2- deoxyglucose (2-DG, Port C) to achieve concentrations of 1 mM, 2 μM, and 50 mM, respectively, after injection. During the test 10 mM glucose, 5 μM oligomycin and 100 mM 2-deoxyglucose (2-DG) (Sigma-Aldrich) were injected to each well after 18, 36 and 65 minutes respectively. After injections, measurements of 2 minutes were performed in triplo, preceded with 4 minutes of mixture time. The first measurements after oligomycin injections were preceded with 5 minutes mixture time, followed by 8 minutes waiting time for the mitochondrial stress test and 5 minutes mixture time followed by 10 minutes waiting time for the glycolysis stress test.

For OXPHOS assay, cells were washed and preincubated with basic medium contained 1 mM D-glucose and 2mM glutamine. The sensor cartridge was loaded with oligomycin (Port A), FCCP (Port B), and rotenone/antimycin A (Port C) to achieve final concentrations of 2 μM, 1 μM, and 2μM, respectively, after injection. Oxygen consumption rate (OCR) was measured using the measurement protocol described above. Both ECAR and OCR were normalized to individual protein amount, and data was analysed using the XF Mito Stress Test Report Generator.

### Clinical specimens

Skin specimens were obtained from the Guangdong Provincial Hospital of Chinese Medicine (China) with Institutional Review Board approval (BF2019-108-01), and written informed consent was obtained from all patients. A total of 10 cases, including 5 cases of psoriasis patients and 5 cases of healthy people with an age ranging from 23 to 46 y (mean, 37 y) and 31 to 49 y (median, 38 y) respectively, were enrolled in this study. Samples were formalin-fixed paraffin-embedded and were furtherly used for in situ hybridization detection of miR-31 and immunohistochemistry staining of specific proteins.

### Animal experiments

An imiquimod-induced mouse model (IMQ) of psoriasis was introduced in this study. 8-week-old BALB/c mice were obtained from Guangdong experimental animal center, follow with shaved and chemically depilated dorsal hair. In brief, to establish an IMQ model, 50mg of imiquimod cream (5%) (Med-Shine Pharmaceutical, China) was applied topically onto the dorsal skin of mice continuously for 6 days. In this study, two independent animal experiments were processed. In order to investigate metabolic enzymes expression in association with the pathology development of psoriasis, we harvested lesioned skin from IMQ mice (n=3) every other day, from day 0 to day 10. In addition, mouse experiment were performed in parallel to study the therapy effect of CB-839 in psoriasis. Starting from the second day of imiquimod induction, BALB/c mice were assigned to received DMSO, CB839 (0.6 mg/mouse/day), or Calcipotriol and Betamethasone Dipropionate Gel (50 μg/mouse/day, commercial name: Xamiol, LEO laboratories Lsd., USA) topically prior to imiquimod treatment in a total of 6 days. The mouse experiment was approved by Animal ethics committee of Guangdong Provincial Hospital of Chinese Medicine (China, No. 2018068).

### Immunohistochemistry (IHC) and Hematoxylin & Eosin Y staining (HE)

Biopsies were harvested from dorsal skin of mice and fixed with 10% neutral buffered formalin (NBF) about 48h. After paraffin-embedding, all samples were cut into 4μm sections and attached to microslide. Heat-induced antigen repair was performed by using a pressure cooker and natural cooled down about 10 mins) in. Sections were deparaffinized, heat retrieved (buffer with 1 citrate buffer, pH 6, cooked for 2 mins in pressure cooker and kept in 94–96 °C for 10min, cooled naturally), perforated (0.2% TBST, 10 min), blocked in 2% BSA (Sigma, A9418) and then incubated with antibodies listed in Extended table 7. The immunostaining was performed using an EliVision System-HRP DAB (MXB, DAB-2031). In the end, sections were counterstained with hematoxylin, dehydrated and cover slipped. HE was performed under the manufacturer’s instructions. The staining was quantified using Image J. All IHC and HE staining shown are representative of three or more sections.

### In situ hybridization

For miR-31 in situ hybridizations, digoxigenin (DIG)-labelled probes (Exiqon) were used following the manufacturer’s protocol. Both DIG-labelled miR-31 and scrambled probes (Exiqon) were hybridized at 61°C. U6 probe was used as the positive control. In situ signals were detected by staining with Anti-DIG-AP antibody (Sigma, cat.11093274910) and developed using NBT/CBIP purple substrate (Sigma, cat.11697471001). Sections were counterstained with nuclear fast red (VECTOR, cat. H3403) for 1 min and washed by flowing water. The quantification was performed by using Image J.

### Targeted analysis of the cellular metabolome

The targeted metabolomics was performed using Hydrophilic Interaction Chromatography - Mass Spectrometry (HILIC-MS) in negative mode. Cells washed with PBS were transferred to 2 mL Eppendorf tubes and homogenized in ice cold 80% methanol. After centrifugation at 15000g for 10 min the supernatant was transferred to a clean Eppendorf tube and evaporated to dryness at 35°C in a Pierce Reacti-Therm III under a gentle stream of nitrogen gass. The residue was dissolved in 70 µL 5% acetonitrile in milli-Q water, centrifugated for 5 min at 10000g, and transferred to a clean injection valve for UPLC-MS analysis. The UPLC–MS system was composed of a Waters Acquity M-class coupled to a Waters VION Q-ToF. Chromatographic separation was performed using a SeQuant ZIC-cHILIC column (150x1 mm, 3 µm) operated at 70 µL min-1. Autosampler temperature was kept at 4°C, the column oven was kept at 40°C. Mobile phase A consisted of 95% acetonitrile in Milli-Q water containing 5 mM ammonium acetate (pH6.8). Mobile phase B consisted of 5% acetonitrile in Milli-Q water containing ammonium acetate (pH6.8). The step gradient program took 30 min. The gradient was kept at 25% B for 2 min after an injection of 5 µL of sample. Thereafter the gradient linearly increased to 70%B in the following 3 min and was kept at 70% for 4 min. The gradient linearly decreased to 25% B in 2 min and the column was allowed to recondition for 19 min prior to the next analysis. MS analysis was performed using a VION Q-ToF operated in full scan mode (75-900 m/z) and in negative mode only. Capillary voltage was set to 1 kV, the sampling cone was set to 20V. Source temperature was 120°C, desolvation temperature was 450°C. Cone gas flow was 50 L h-1 and desolvation gas flow was set to 800L h-1. The lock mass was Leucine enkephaline. Standard solutions were used to determine both retention time and m/z values (△ m/z < 2 ppm). Analysis was conducted with the system operation software UNIFY.

### Amino acid assay

A volume of 15 µL medium was transferred to a labelled 1.5 mL Eppendorf vial. A volume of 285 µL 80% acetonitrile also containing internal standards (final concentration 10 µM) was added and the sample was thoroughly mixed. The sample was centrifuged for 10 min at 17000xg in an Eppendorf centrifuge. A volume of 250 µL was transferred to a new, labelled Eppendorf vial and evaporated to dryness. Cell samples harvested in 300 µL ice-cold methanol were subjected to the same protol. The residue was dissolved in 70 µL of borate buffer (pH 8.2) by thorough mixing and the derivatisation was started by adding 20 µL of AccQ-Tag reagent solution prepared according to the suppliers protocol (Waters, Etten-Leur, The Netherlands) after which the samples were vortex mixed and incubated at 55°C for 10 min. The sample was evaporated to dryness and the residue was dissolved in 120 µL 10% acetonitrile containing 0.1 mM formic acid and transferred to an LC sample vial.

Analysis was performed on a system consisting of an Ultimate 3000 LC and an LTQ-Orbitrap XL (Thermo Scientific, Breda, The Netherlands). As a column a Waters HSS T3 (2.1x100 mm, 1.8 µm) was used, kept at a temperature of 40°C in the column oven. Eluent A used for analysis was milliQ water containing 0.1% formic acid, eluent B consisted of acetonitrile containing 0.1% formic acid. The LC gradient used for separation commenced by injecting 1 µL of sample and started at 0% B for 5 min. followed by a 10 min linear gradient to 75% B. In 0.5 min the gradient increase to 100% B and kept there for 1 min before returning to 0% B. The column was allowed to regenerate for 2.5 min prior to a next analysis. Total runtime was 18 min; flow rate was 400 µL/min. The mass spectrometer was operated in ESI-positive mode, full scan 100 – 1000 m/z, capillary temperature 300°C, sheath gas: 35, aux gas:2, resolution 30000, capillary voltage 3 kV.

### Glutamine tracing

HaCAT cells were transfected with control scrambled miRNA or miR-31. After 32 hrs culture medium was replaced with 0.5%FBS DMEM with 1mM glucose, without glutamine for 16h. To start glutamine labelling, the medium was replaced with DMEM contains 0.5%FBS, 1mM glucose, and 2mM 13C-glutamine. Samples were harvested at the indicated time points after start of glutamine labelling. Medium was collected and snap frozen and cell were immediately harvested in 100% ice cold methanol

Analysis was performed as described (Ciapaite *et al*, 2020), with following modifications. Prior to analysis, calibration samples were prepared by dilution and addition of internal standards. Calibration samples were prepared in the concentration range from 0.08-80 µM. 20 µl of Internal standard mix (100 µM) was added to e500 µl methanol sample extract or 50 µl standard and evaporated under a gentle flow of nitrogen at 40 °C. To the dried extract 25 µl 0.1% NaOH and 25 µl 10 mg/ml O-(2,3,4,5,6-pentafluorobenzyl)hydroxylamine (PFBHA) in Milli-Q water, was added, vortexed for homogenization and for derivatization incubated for 30 min at 20°C. For determination of extracellular metabolites, to 50 µl cell medium, 20 µl of Internal standard mix (100 µM), 25 µl 0.1% NaOH and 25 µl 10 mg/ml O-(2,3,4,5,6-pentafluorobenzyl)hydroxylamine (PFBHA) in Milli-Q water is added, vortex and derivatize 30 min at 20°C.

The derivatized metabolites were separated on a Sunshell RP-Aqua column (150 m× 3 mm i.d., 2.6 μm; ChromaNik Technologies Inc., Osaka, Japan). The following eluents were used: solvent A: 0.1% formic acid in H2O (v/v); solvent B: 0.1% formic acid in ACN (v/v). Gradient elution was as follows: 0–2.75 min. isocratic 0% B, 2.75–3.5 min. linear from 0% to 70% B, 3.5–6.0 min. isocratic 70% B, 6.0–6.2 min. linear from 70% to 100% B, 6.2–9.0 min. isocratic 100% B, and 9.0-9.2 min. linear from 100% to 0% B, with 9.2-12 min. for initial conditions of 0% B for column equilibration. Settings used were: Flow rate: 0.6 ml/minColumn temperature: 40 °C; Autosampler temperature: 15 °C; Injection volume: 5 µl.

The eluent was analyzed on a Q-Exactive HF mass spectrometer with the following settings: detection mode: Full scan in negative ionisation mode; scan range: 70 to 900 m/z; resolution: 240000; AGC target: 1e6; Maximum IT: 200 ms; capillary voltage: 4kV; capillary temperature: 300 °C; sheath gas: 50; aux gas: 2; spare gas: 0; S-lens RF level: 65; Ion source: ESI.

### Analysis of cytokine production by multiplexed particle-based flow cytometric assay (Luminex)

Cell culture supernatants were collected, stored at −80°C and processed as described(Scholman *et al*, 2018). Cytokine concentrations were measured with the Bio-Plex system in combination with the Bio-Plex Manager software, version 4.0 (Bio-Rad Laboratories, Hercules, CA, USA), which employs the Luminex xMAP technology(Scholman *et al*., 2018).

### Induction of *in vitro* Th17 differentiation

Spleens were harvested from C57BL/6 mice for lymphocyte isolation following the protocol of Bedoya (Bedoya *et al*, 2013). CD4+CD25- naive T cells were subsequently isolated by negative selection using magnetic beads (Cat: 130-104-453, Miltenyi Biotech, Germany). Naive CD4+ T cells were stimulated with plate-bound anti-CD3 (LOT: 4310627, eBioscience) and 1.5 μg/ml soluble CD28 monoclonal antibodies (Cat: 557393, BD biosciences pharmingen) in DMEM medium and cytokines (IL-6, 20 ng/ml; TGF-b1, 5 ng/ml; IL-21, 25 ng/ml) or metabolites (Glutamate 25μM; Aspartate 25μM) for a period of 5 days, at which point cells were collected and were stained with the following antibodies: CD4-PE (LOT:2004774, Invitrogen), RORγt-APC (LOT:2279171, Invitrogen) and CD25-PE-Cy7(LOT:2073783, Invitrogen) and were analysised on a Novo Quanteon (Aglient).

---

**Extended data Figure 1.**
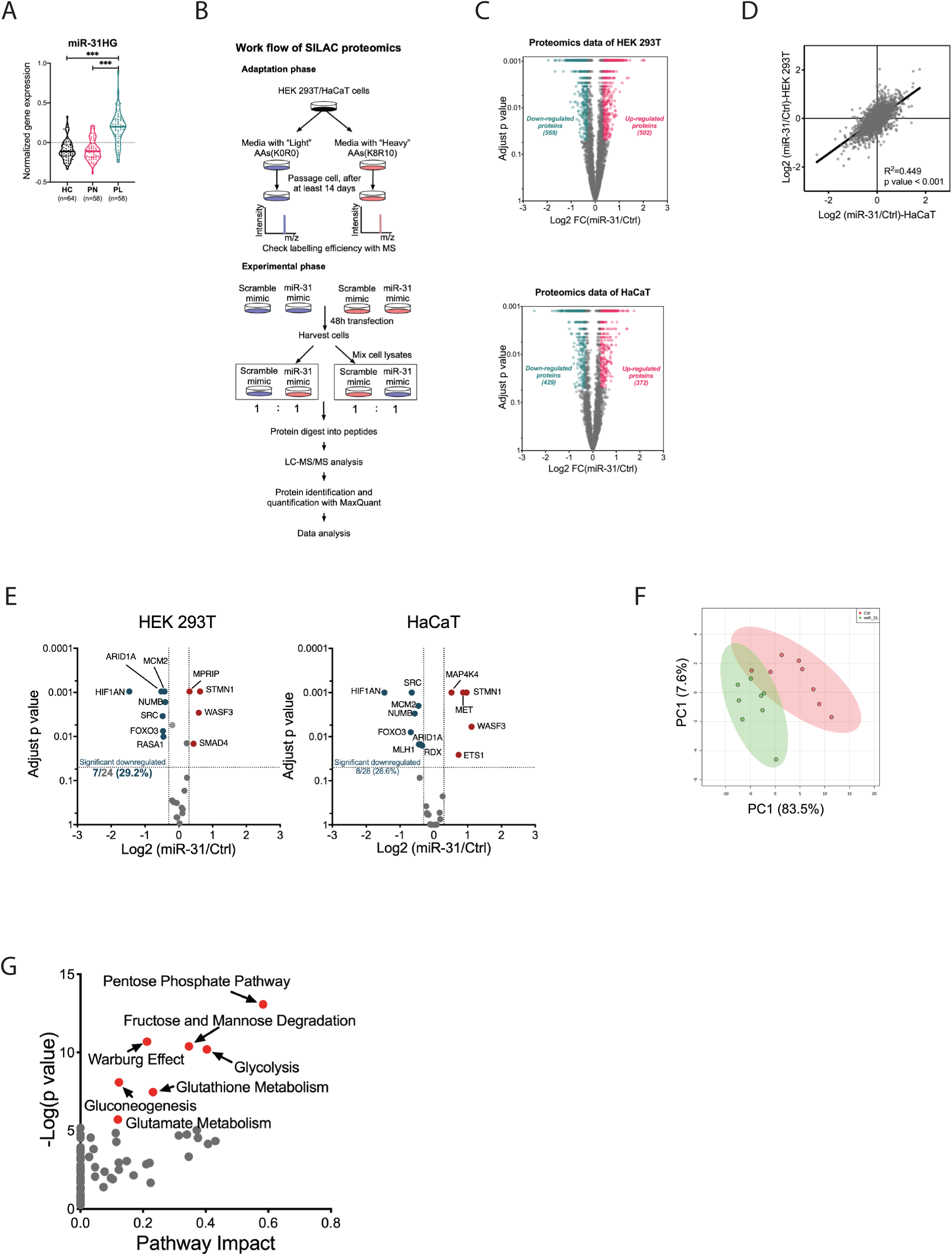
A. Violin plot of miR-31 host gene (miR-31 HG) expression in human skin biopsies. Data extracted from an online microarray study (GSE13355). n=64/58 in healthy control (HC)/psoriasis patient (PN, psoriatic non-lesional skin; PL, psoriatic lesional skin), respectively. mean ± SD, p values were calculated by One-way ANOVA with Sidak test, ***, p<0.001. B. Schematic representation of the workflow for SILAC proteomics. C. Volcano plots summarizing SILAC proteomics data in HEK 293T (upper panel) and HaCaT (bottom panel) cell lines. D. Correlation analysis of SILAC proteomics data of two cell lines. E. Volcano plot representations of changes of validated targets of miR-31 in HEK 293T cell (left) and HaCaT cell (right) proteomics. F. Principal component analysis (PCA) shows a clear distinction between the miR-31 and normal HEK 293T cells, indicating a specific metabolism signature induced by miR-31 overexpression. G. Metabolic pathway enrichment analysis using miR-31-induced significant changed metabolites as input. Those most predicted pathways are highlighted in red.

**Extended data Figure 2.**
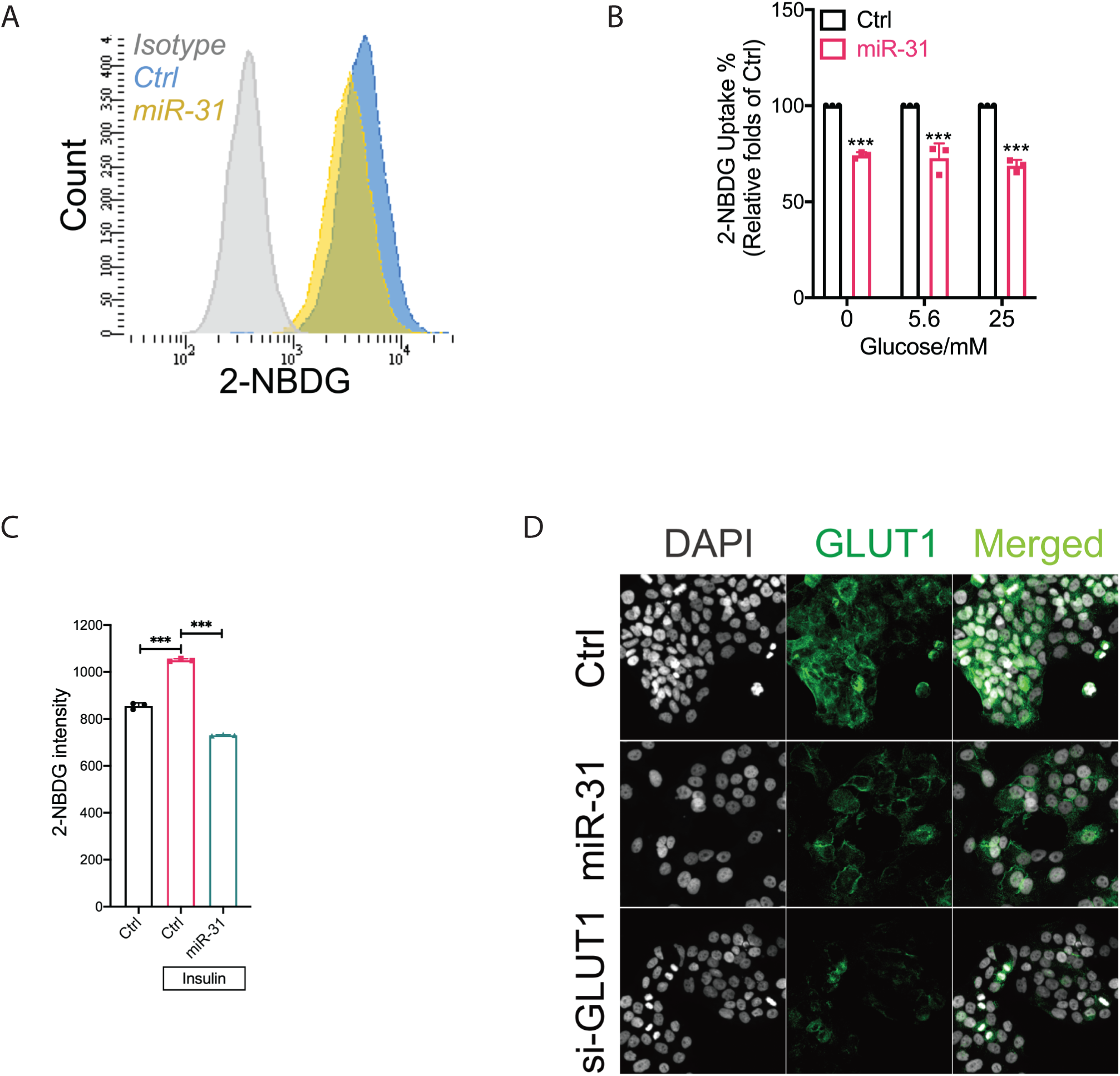
A. Measurement of 2-NBDG uptake in HaCaT cells by flow cytometry. B-C. 2-NBDG uptake of HaCaT cells in response to glucose (B) and insulin (C) additions. n=3, mean ± SD, ***, p<0.001. D. Immunostaining of GLUT1 in HaCaT cells of 48 hours treatments of scramble, miR-31 mimic and GLUT1 siRNA (si-GLUT1). DAPI is grey and GLUT1 is green. Similar results were obtained from three independent experiments. p values (indicated about relevant comparison) were calculated by Student’s t-test and One-way ANOVA with Sidak test (C), Two-way ANOVA with Sidak test (B).

**Extended data Figure 3.**
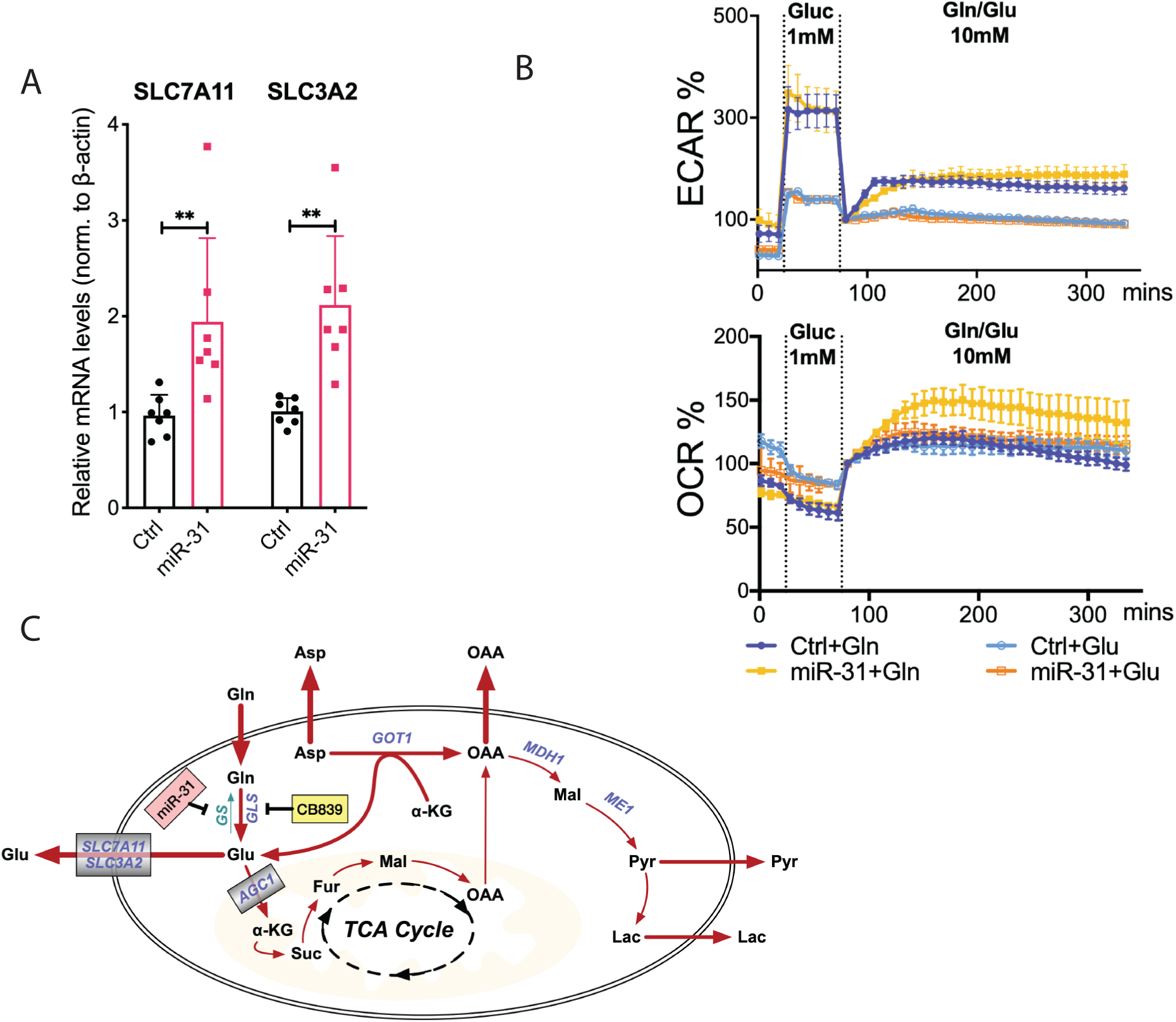
A. qRT-PCR analysis of SLC7A11 and SLC3A2 in HaCaT cells. n=7, mean ± SD, p values was calculated by One-way ANOVA with Sidak test, *** means p<0.001. B. Baselined ECAR and OCR of HaCaT cells in response to glutamine or glutamate addition in a long-term observation. n=5, mean ± SD. C. A summary scheme showing how miR-31 rewires glutamine metabolism.

**Extended data Figure 4.**
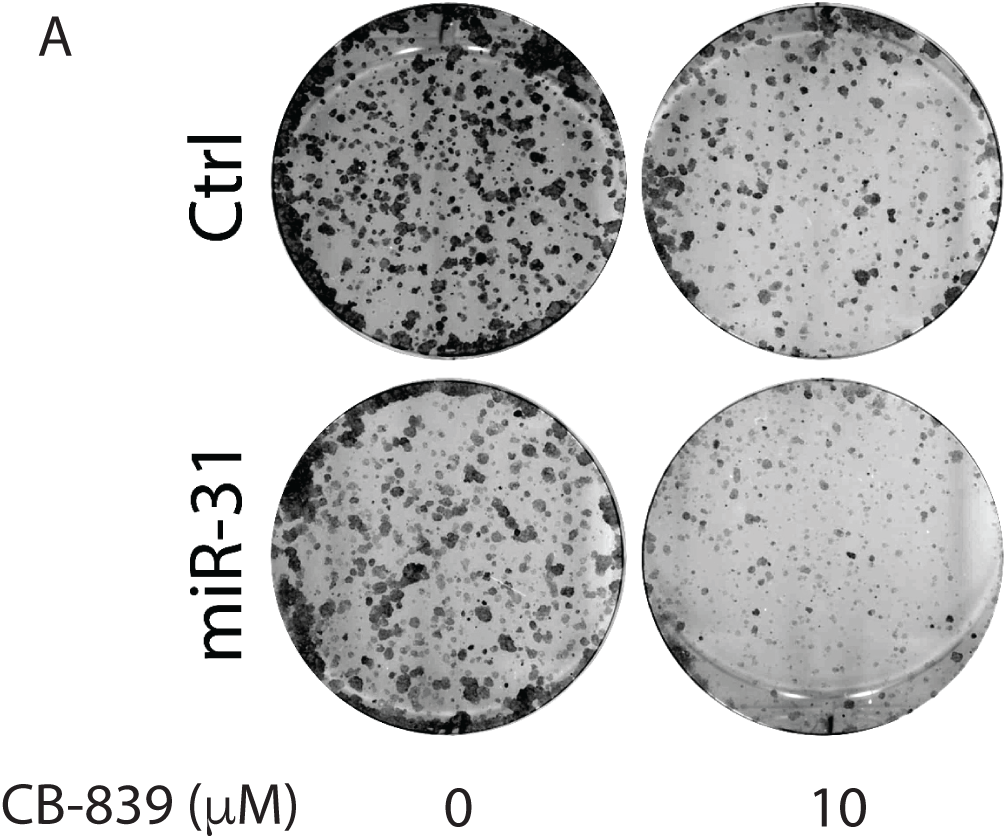
A. Representative picture of cell colony forming assay of HaCaT cells in response to CB839 treatment, n=3 replicates.

**Extended data Figure 5.**
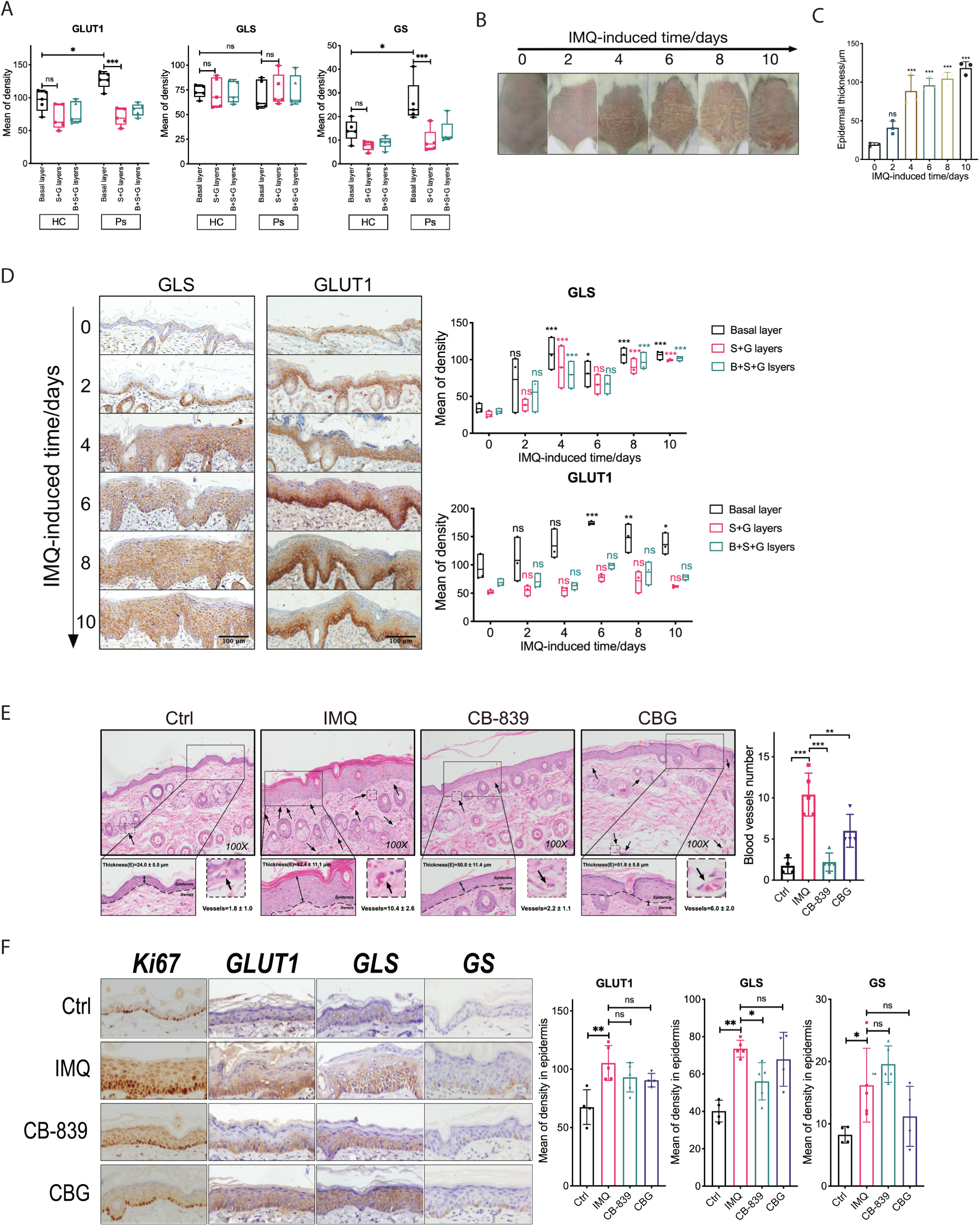
A. Quantification of results of GLUT1, GLS, and GS staining of human biopsies. S+G layers, spinous layer and granular layer of epidermis; B+S+G layers, basal layer, spinous layer and granular layer. *, p<0.05; ***, p<0.001. B. Representative pictures of skin from imiquimod (IMQ)-induced mice in a time course, n=3. C. Quantification of skin epidermal thickness of mice with IMQ induction in a time course, n=3. D. GLS and GLUT1 staining of skin from IMQ-induced mouse in a time course. Quantification results showed on right panel. n=3, *, p<0.05; **, p<0.01; ***, p<0.001. E. Histological staining of skin indicating lesser blood vessels in CB-839 treatment mice. Blood vessel was highlighted by arrow. Quantification on right panel, n=4-5, mean ± SD, **, p<0.01; ***, p<0.001. F. Ki67, GLUT1, GLS, and GS staining of skin from mice with different treatments. Quantification results on right panel, n=4-5, mean ± SD, *, p<0.05; **, p<0.01. p values (indicated about relevant comparison) were calculated by One-way ANOVA with Sidak test (C, E and F), or Two-way ANOVA with Sidak test (A and D).

**Extended Table 1:**
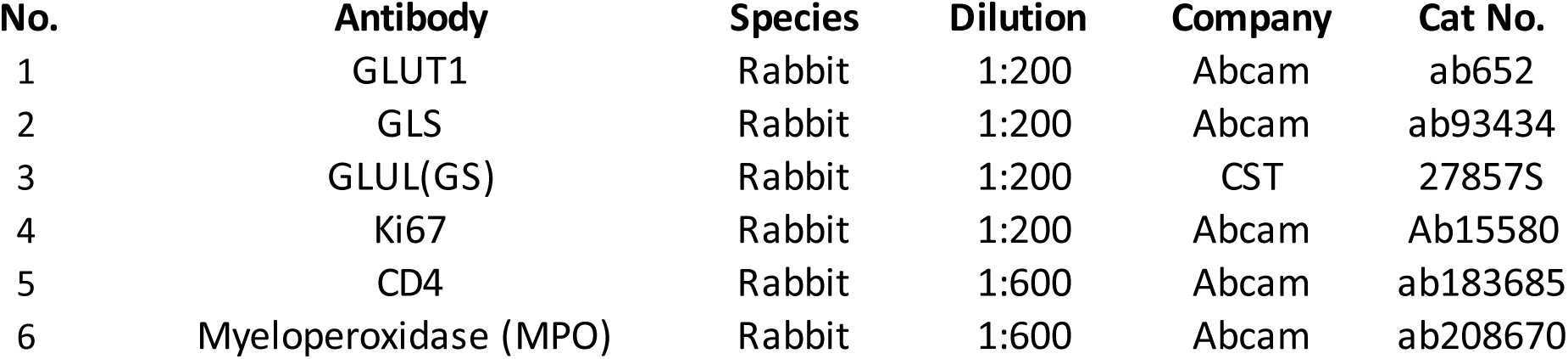
listing of overlap between proteomic changes and literature validated miR-31 targets.

**Extended Table 2:**
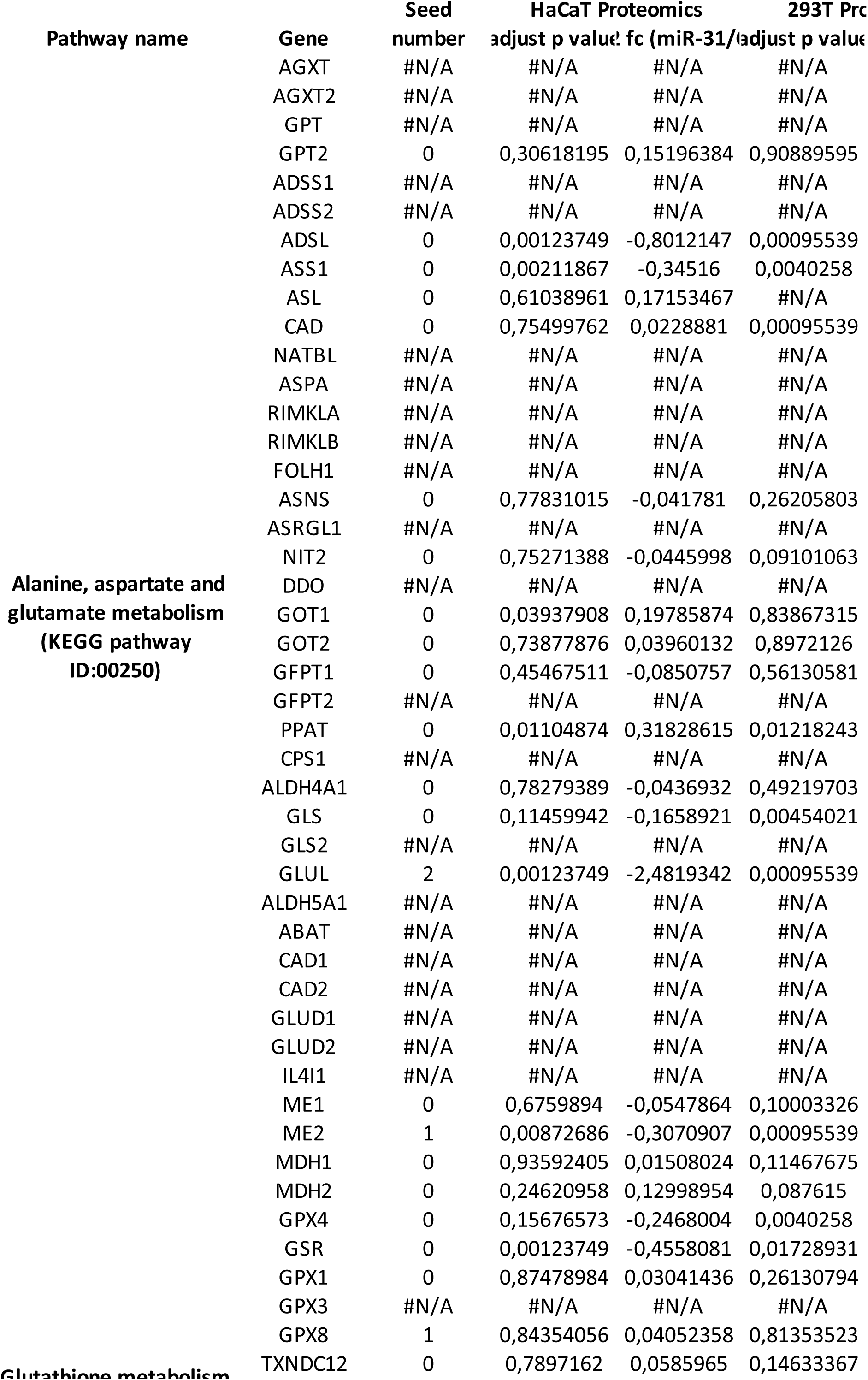
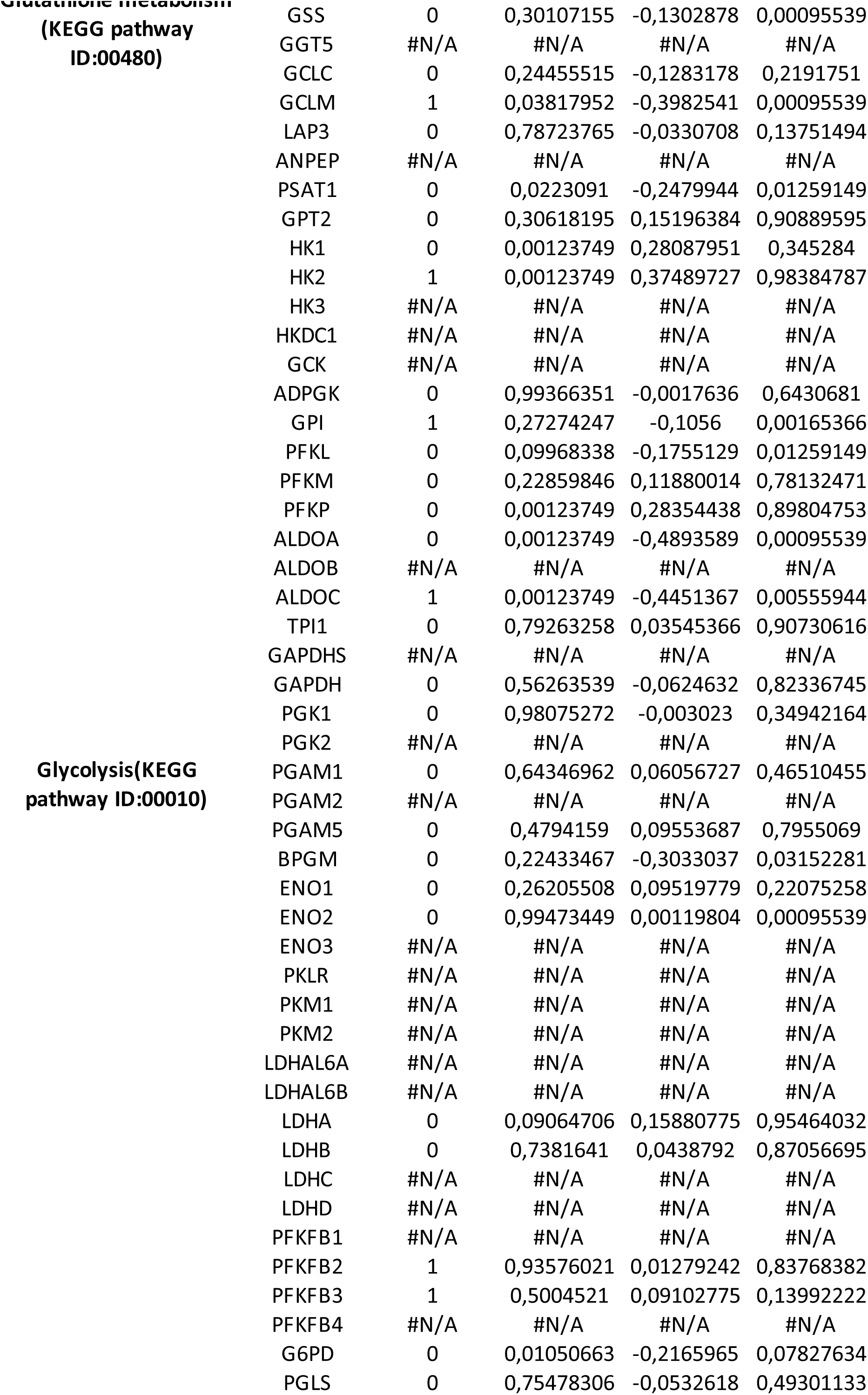
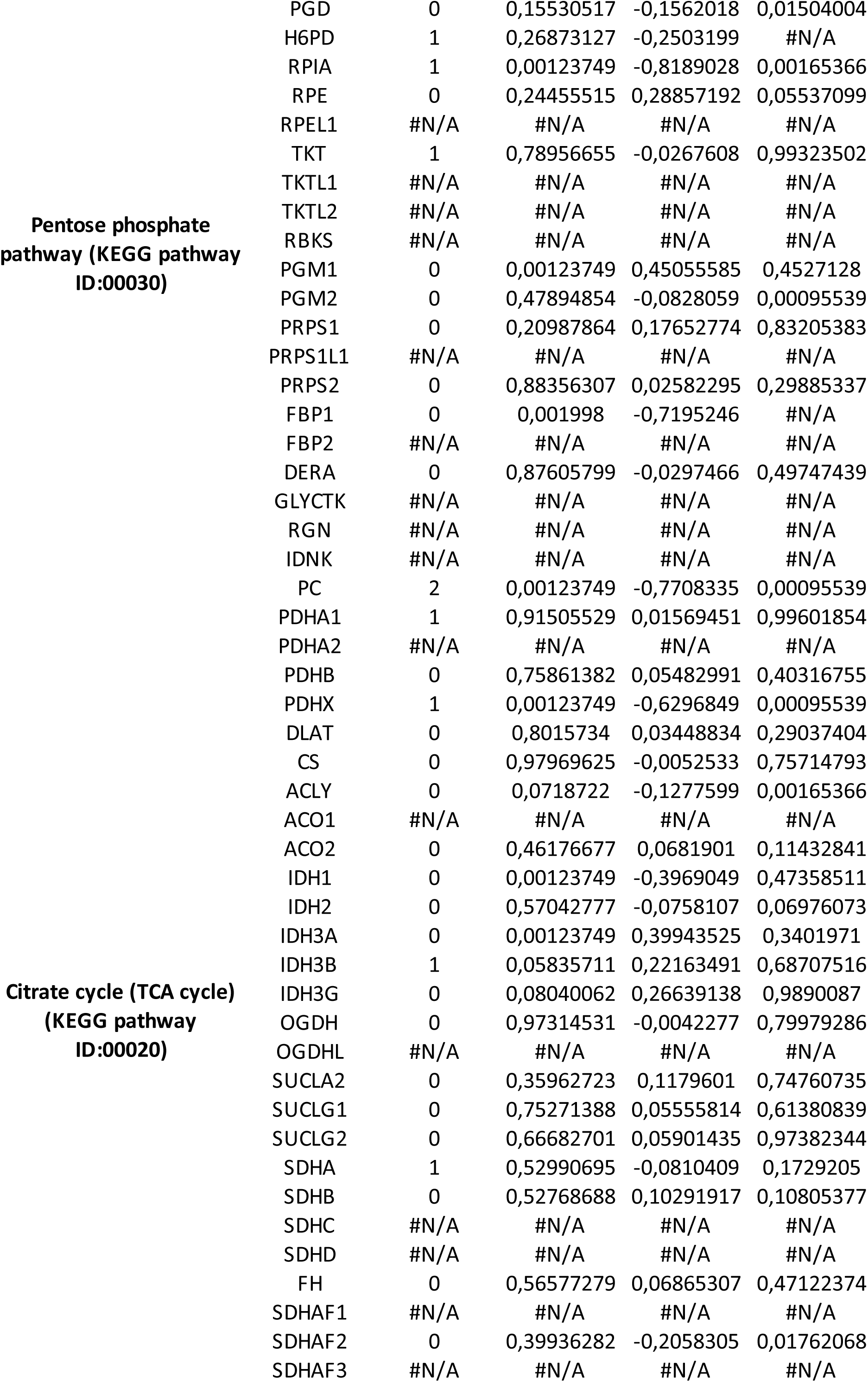
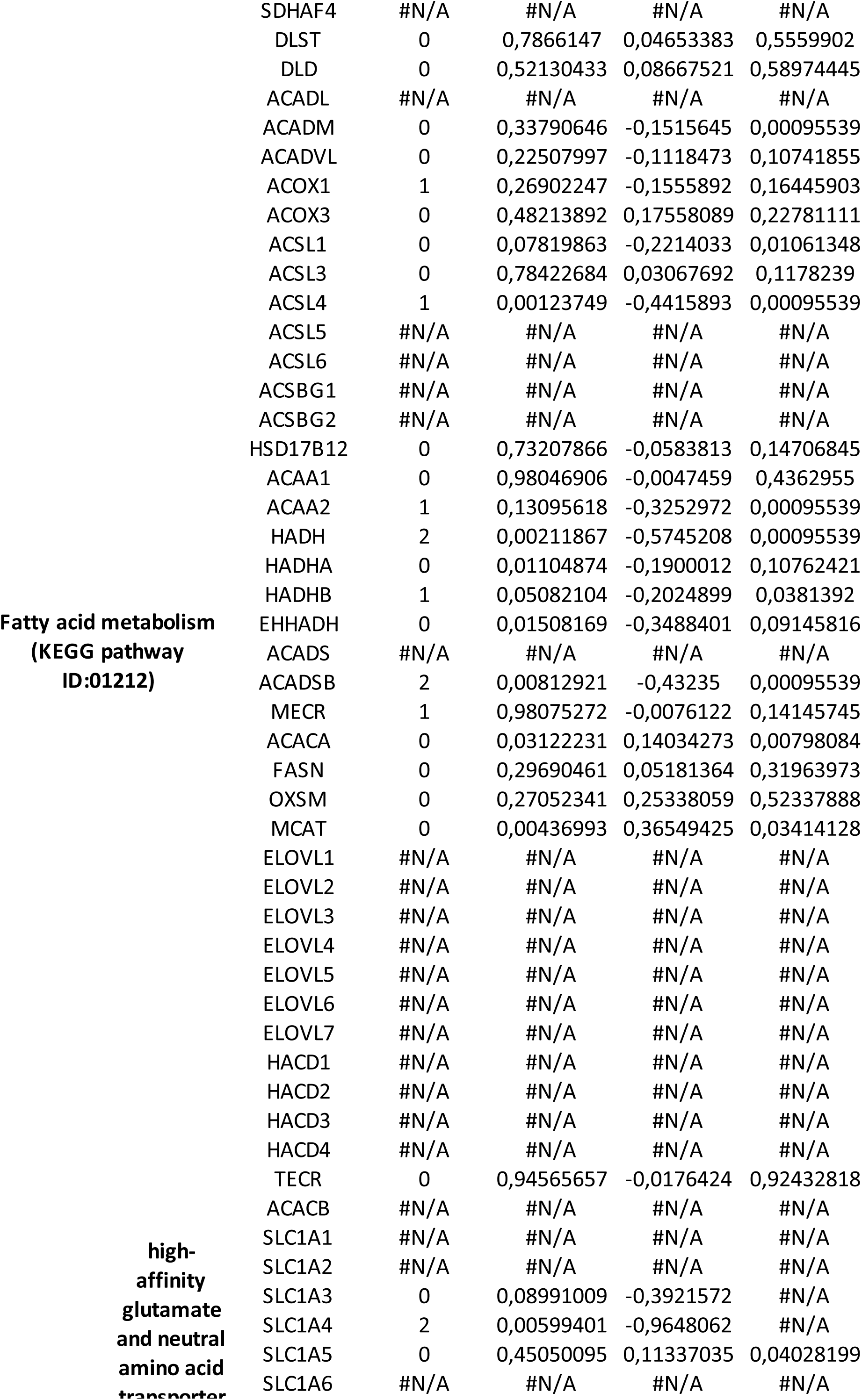
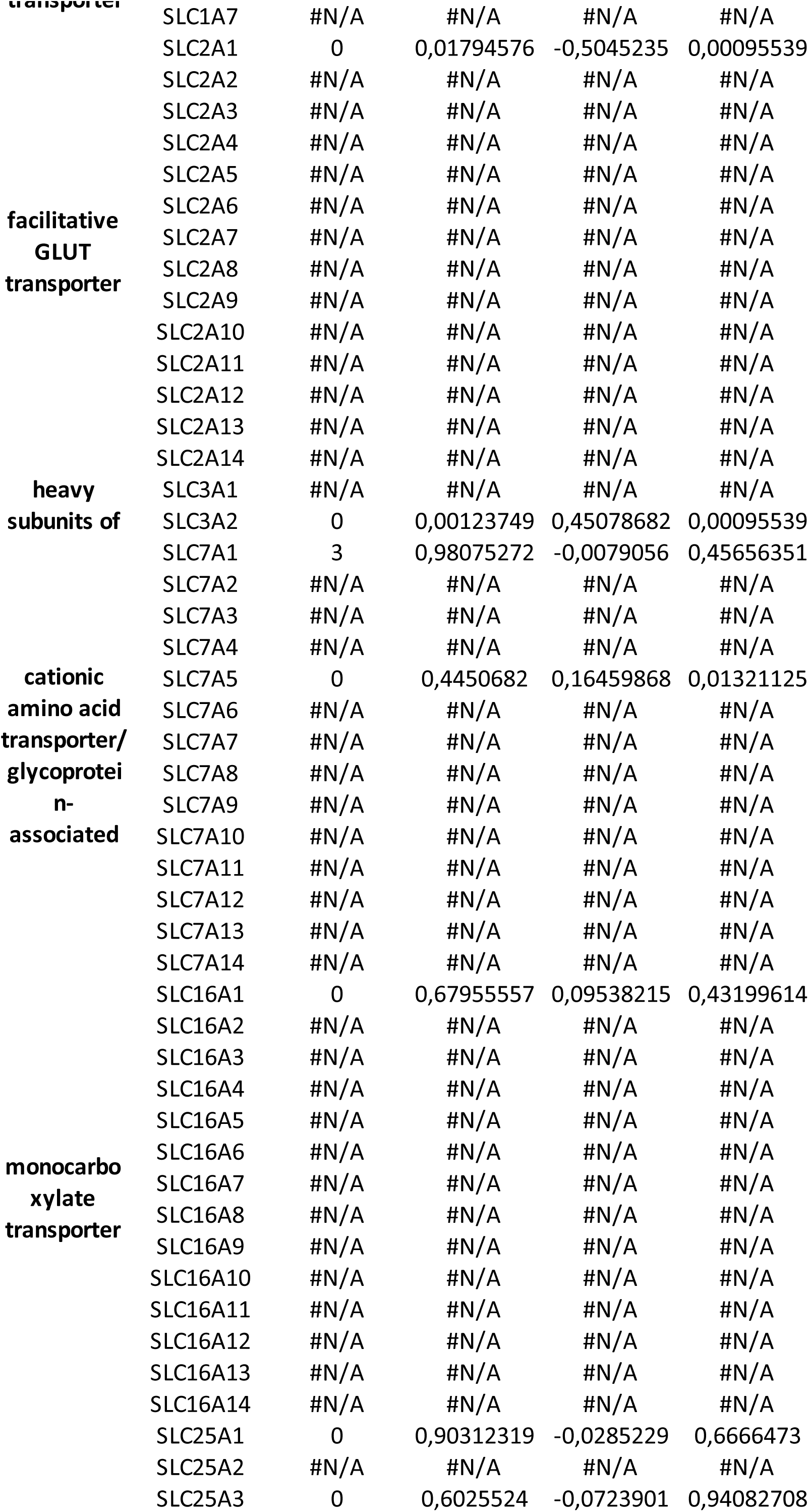
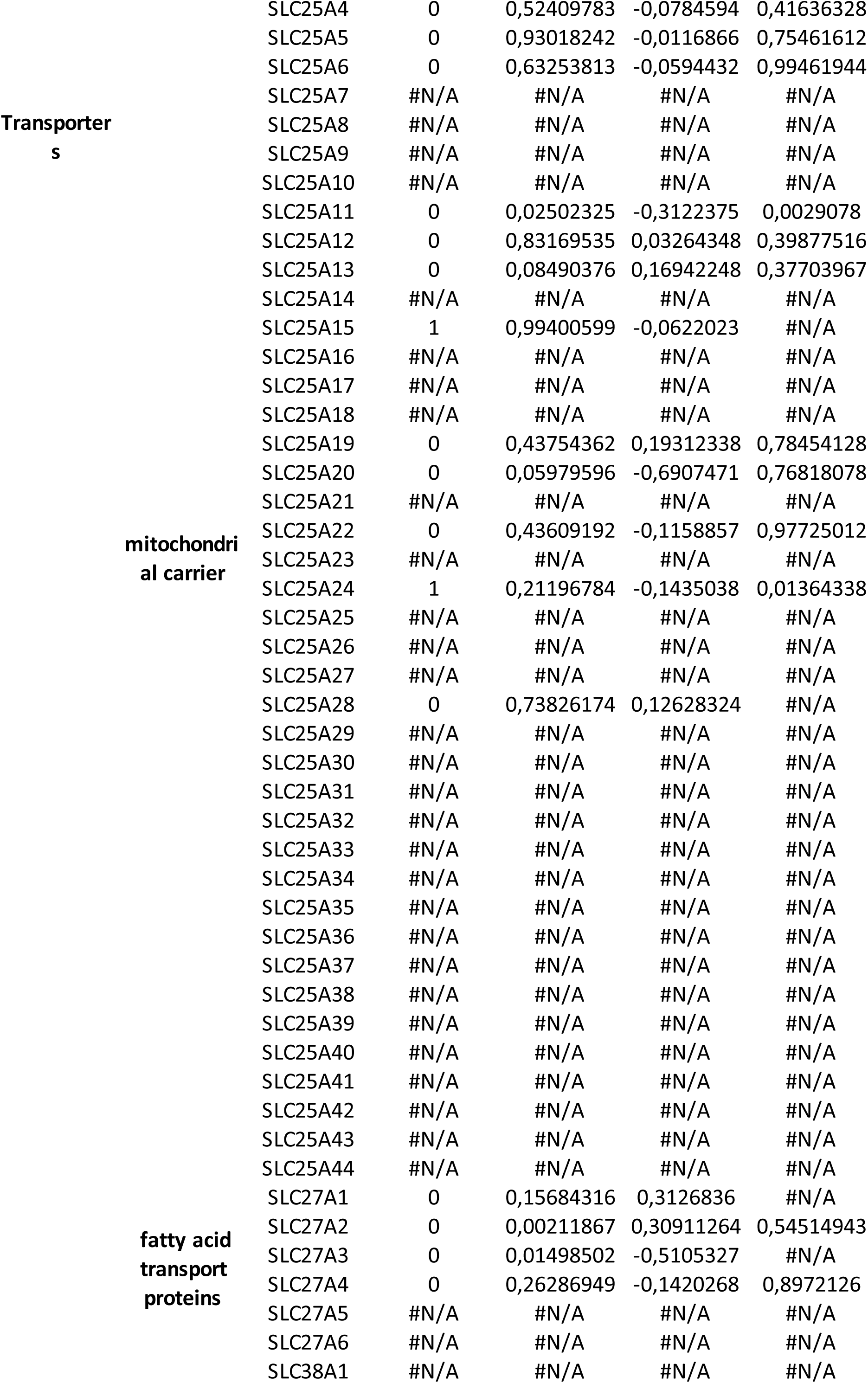
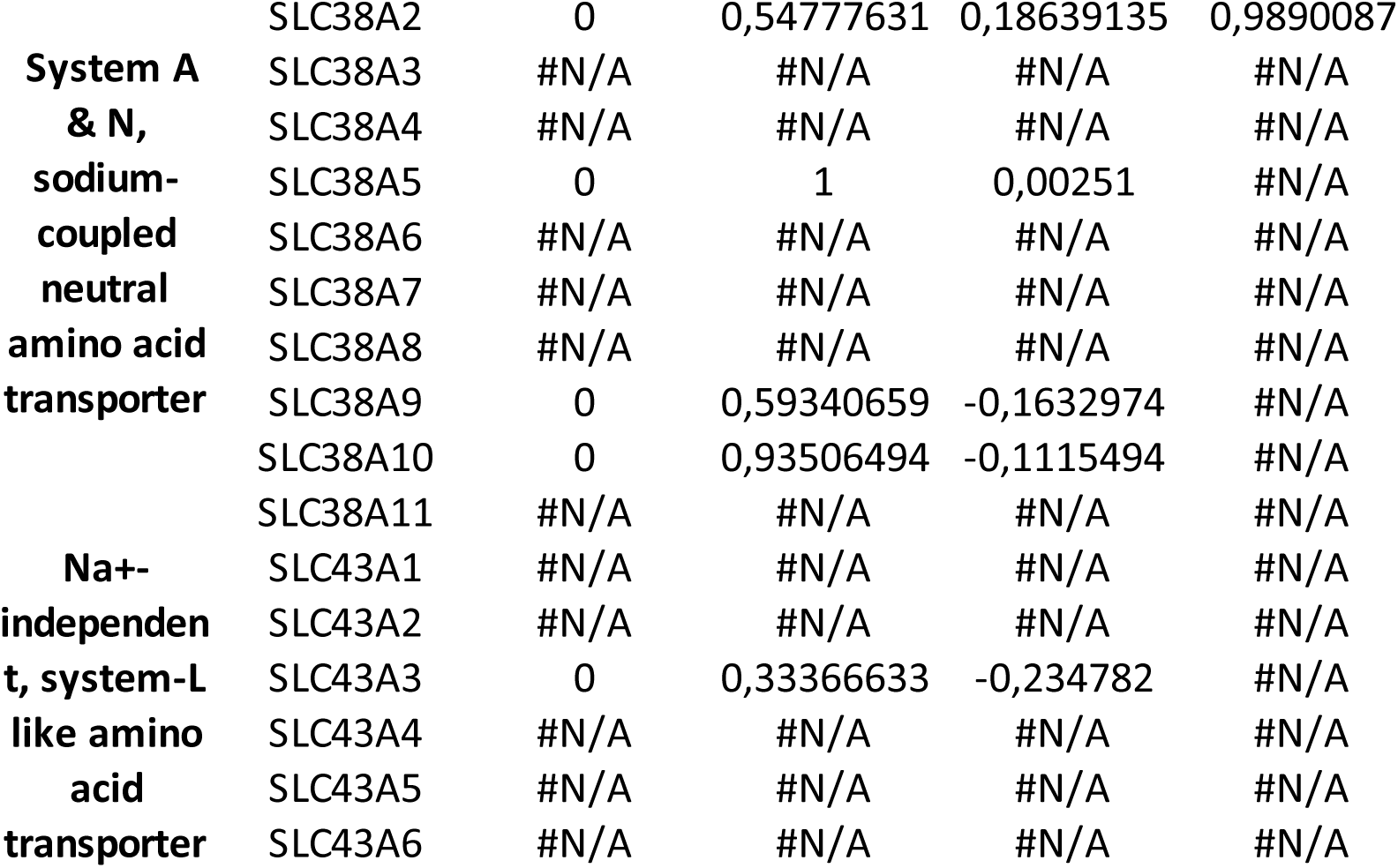
miR-31 seed sequences in 3’UTR of metabolic genes

**Extended Table 3:**
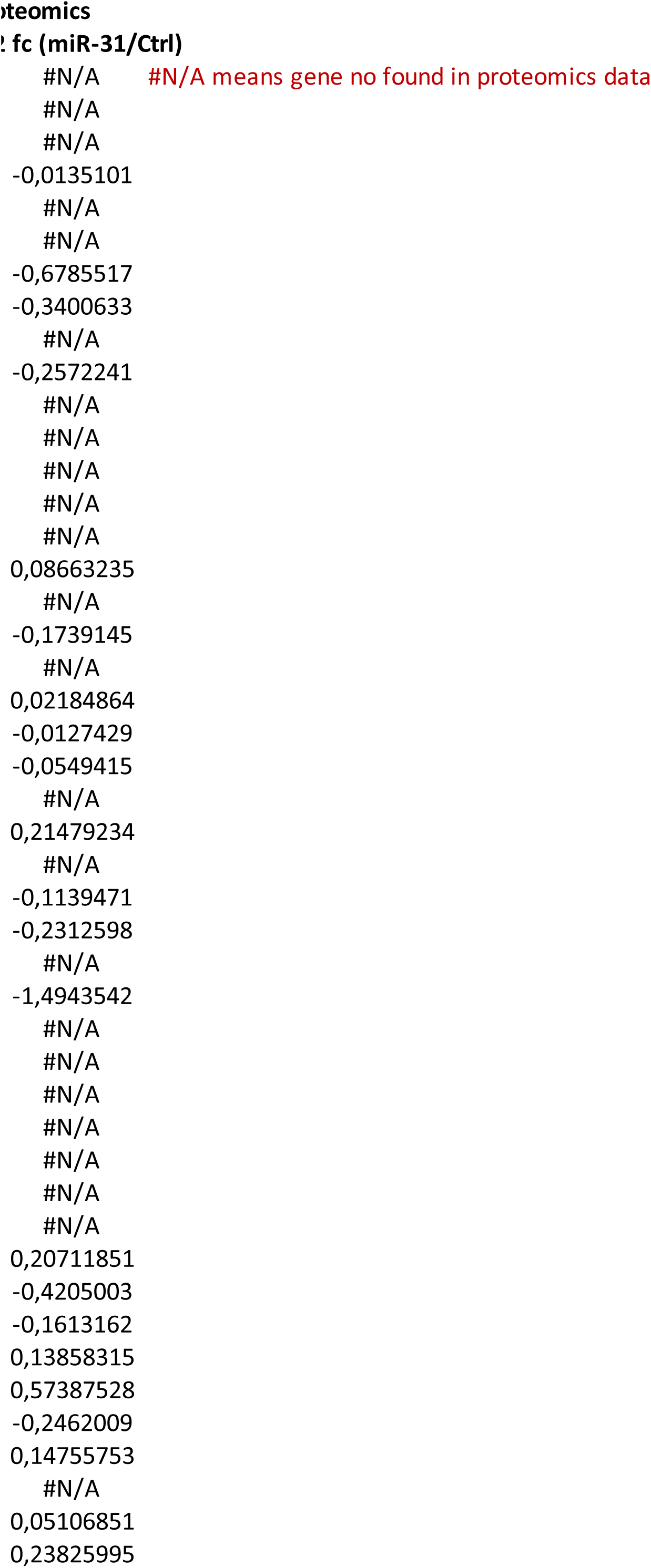
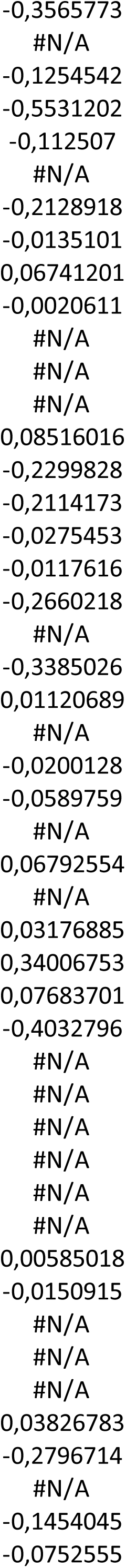
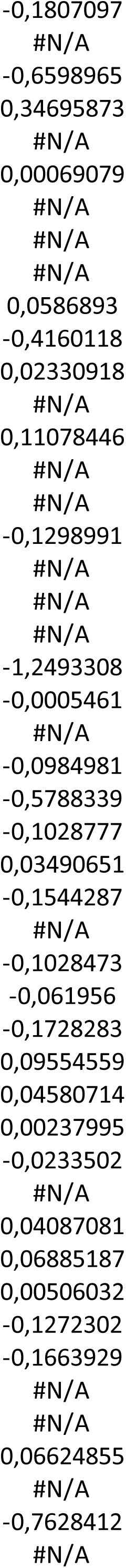
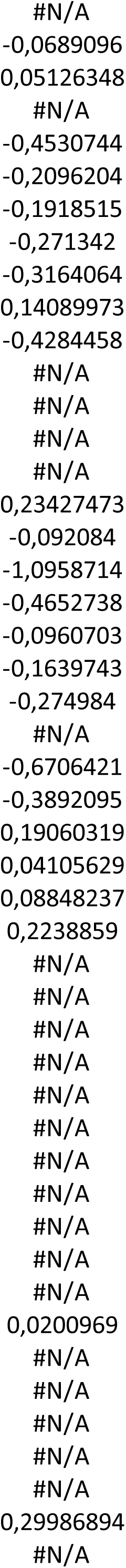
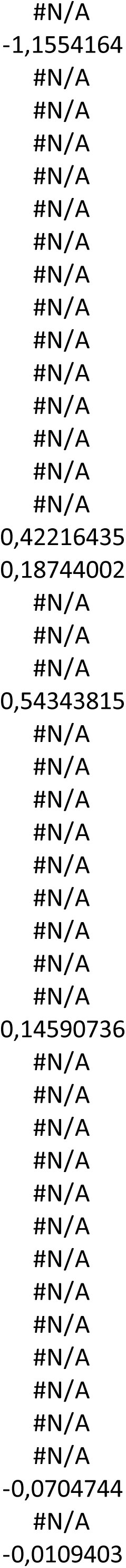
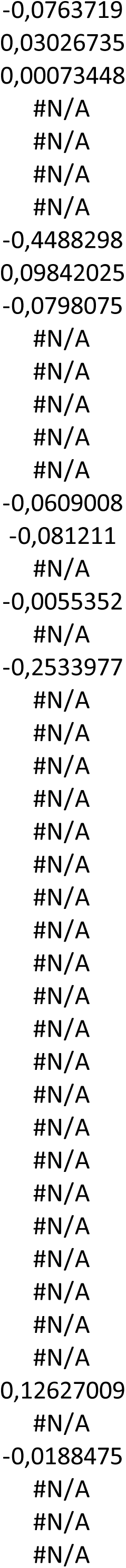
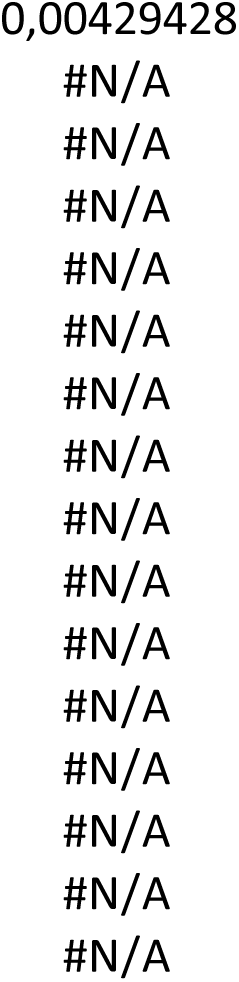
metabolite profiling upon miR-31 overexpression (targeted metabolomics, Fig.1d)

**Extended Table 4.** is: cytokine production in response to miR-31 over expression and CB-839 treatment. (belonging to Luminex data, Fig. 4e)

| SAMPLES<br>Group | Ctrl_1<br>Ctrl | Ctrl_2<br>Ctrl | Ctrl_3<br>Ctrl | Ctrl_4<br>Ctrl | Ctrl_5<br>Ctrl | Ctrl_6<br>Ctrl |
| --- | --- | --- | --- | --- | --- | --- |
| GSSG | 0,355 | 0,269 | 0,371 | 0,346 | 0,342 | 0,392 |
| GSH | 5,769 | 7,449 | 6,763 | 8,4 | 9,108 | 7,071 |
| NAD | 0,218 | 0,214 | 0,269 | 0,251 | 0,247 | 0,302 |
| NADH | 0 | 0 | 0 | 0 | 0 | 0 |
| NADP | 0,006 | 0,007 | 0,01 | 0,012 | 0,011 | 0,012 |
| NADPH | 0,001 | 0,002 | 0,002 | 0,002 | 0,001 | 0,002 |
| FAD | 0 | 0,001 | 0,001 | 0,001 | 0,001 | 0,002 |
| Glucose | 0,948 | 0,982 | 1,399 | 1,305 | 1,413 | 1,314 |
| Glucose_6P | 1,933 | 1,848 | 2,747 | 2,125 | 2,644 | 3,31 |
| Fructose_6P | 2,088 | 2,037 | 3,012 | 2,474 | 3,088 | 3,436 |
| Fructose_1_6P | 1,171 | 3,269 | 6,21 | 5,43 | 6,057 | 5,854 |
| Glyceraldehy | 0,331 | 0,348 | 0,299 | 0,349 | 0,37 | 0,476 |
| Lactic_acid | 0,022 | 0,033 | 0,025 | 0,04 | 0,038 | 0,026 |
| Ribulose_5P | 0,209 | 0,226 | 0,328 | 0,277 | 0,354 | 0,354 |
| Ribose_5P | 0,365 | 0,423 | 0,602 | 0,484 | 0,608 | 0,665 |
| Sedoheptulo | 0,478 | 0,51 | 0,778 | 0,577 | 0,719 | 0,744 |
| Erythrose_4P | 0,064 | 0,569 | 0,1 | 0,033 | 0,307 | 0,693 |
| Citric_acid | 0,015 | 0,263 | 0,156 | 0,141 | 0,502 | 0,122 |
| alpha_Ketogl | 0,005 | 0,005 | 0,005 | 0,006 | 0,007 | 0,005 |
| Fumaric_acid | 0,062 | 0,077 | 0,037 | 0,096 | 0,094 | 0,069 |
| Malic_acid | 0,42 | 0,501 | 0,232 | 0,593 | 0,569 | 0,223 |
| N_Acetylglut | 0,025 | 0,028 | 0,027 | 0,031 | 0,036 | 0,028 |
| Myo—inositol | 21,208 | 24,68 | 36,862 | 26,083 | 32,874 | 34,167 |
| Argininosucc | 0,007 | 0,008 | 0,009 | 0,01 | 0,01 | 0,008 |
| Ribose | 0,034 | 0 | 0,05 | 0,04 | 0,067 | 0,05 |
| Succinic_acid | 0,007 | 0,007 | 0,005 | 0,008 | 0,01 | 0,004 |
| Uracil | 0,004 | 0,002 | 0,002 | 0 | 0,002 | 0,004 |
| Thymine | 0,004 | 0,005 | 0,005 | 0,006 | 0,006 | 0,005 |
| Thymidine | 0,022 | 0,018 | 0,021 | 0,017 | 0,019 | 0,025 |
| Guanosine | 0,004 | 0,005 | 0,005 | 0,006 | 0,005 | 0,005 |
| GMP | 0,008 | 0,008 | 0,009 | 0,009 | 0,01 | 0,009 |
| CDP | 0,002 | 0,003 | 0,004 | 0,003 | 0,004 | 0,004 |
| ADP | 0,051 | 0,103 | 0,116 | 0,145 | 0,153 | 0,122 |
| ATP | 0,179 | 0,408 | 0,896 | 0 | 0,644 | 0,957 |
| GDP | 0,001 | 0,003 | 0,016 | 0,007 | 0,002 | 0,018 |
| UDP | 0,014 | 0,017 | 0,009 | 0,029 | 0,029 | 0,01 |
| UMP | 0,025 | 0,029 | 0,037 | 0,04 | 0,041 | 0,035 |
| TDP | 0 | 0 | 0,002 | 0 | 0,002 | 0,002 |
| TTP | 0,003 | 0 | 0,008 | 0,007 | 0 | 0,008 |
| UTP | 0,046 | 0 | 0,193 | 0,189 | 0,146 | 0,226 |
| CTP | 0,003 | 0,001 | 0,027 | 0,018 | 0,001 | 0,021 |
| GTP | 0,002 | 0,007 | 0,01 | 0,011 | 0,009 | 0,008 |
| UDP_NAcGlc | 2,02 | 1,885 | 3,246 | 2,873 | 2,834 | 3,13 |
| Deoxyribose_ | 0,003 | 0,004 | 0,004 | 0,005 | 0,005 | 0,004 |
| Creatine | 0,011 | 0,008 | 0,008 | 0,008 | 0,008 | 0,008 |
| Arginine | 0,002 | 0,029 | 0,017 | 0 | 0,041 | 0,025 |
| Asparagine | 0,025 | 0,003 | 0,005 | 0,002 | 0,001 | 0,019 |
| Tyrosine | 0,329 | 0,344 | 0,416 | 0,429 | 0,428 | 0,455 |
| Alanine | 0,008 | 0,009 | 0,007 | 0,008 | 0,009 | 0,007 |
| Serine | 0,005 | 0,004 | 0,004 | 0,003 | 0,003 | 0,004 |
| Cysteine | 0,004 | 0,003 | 0,005 | 0,005 | 0,005 | 0,005 |
| Lysine | 0,056 | 0,086 | 0,081 | 0,089 | 0,094 | 0,085 |
| Glutamic_ac | 2,559 | 2,886 | 2,895 | 2,995 | 3,106 | 2,681 |
| Aspartic_acid | 0,024 | 0,044 | 0,037 | 0,027 | 0,043 | 0,028 |
| GABA | 0,105 | 0,127 | 0,119 | 0,139 | 0,146 | 0,115 |
| Ornithine | 0,002 | 0,002 | 0,001 | 0,002 | 0,002 | 0,002 |
| Threonine | 0,02 | 0,003 | 0,007 | 0,005 | 0,004 | 0,01 |
| Methionine | 0,017 | 0,02 | 0,021 | 0,023 | 0,021 | 0,023 |
| Valine | 0,087 | 0,098 | 0,105 | 0,122 | 0,114 | 0,108 |
| Proline | 0,17 | 0,199 | 0,201 | 0,217 | 0,214 | 0,202 |
| Cystine | 0,043 | 0,116 | 0,069 | 0,146 | 0,089 | 0,044 |
| Phenylalanine | 0,075 | 0,089 | 0,097 | 0,114 | 0,114 | 0,101 |
| Leucine | 0,053 | 0,062 | 0,063 | 0,078 | 0,077 | 0,062 |
| Tryptophan | 0,011 | 0,016 | 0,015 | 0,018 | 0,017 | 0,016 |
| Isoleucine | 0,122 | 0,1 | 0,128 | 0,102 | 0,085 | 0,114 |
| Glutamine | 2,76 | 2,805 | 3,076 | 3,199 | 2,931 | 2,915 |
| Taurine | 0,352 | 0,434 | 0,434 | 0,476 | 0,479 | 0,438 |
| N—Acetylornithine | 0,002 | 0,003 | 0,002 | 0,003 | 0,002 | 0,002 |
| Acetoacetic_acid | 0,002 | 0,003 | 0,002 | 0,003 | 0,003 | 0,003 |

| Ctrl_7 | Ctrl_8 | miR31_1 | miR31_2 | miR31_3 | miR31_4 | miR31_5 |
| --- | --- | --- | --- | --- | --- | --- |
| Ctrl | Ctrl | miR_31 | miR_31 | miR_31 | miR_31 | miR_31 |
| 0,339 | 0,351 | 0,411 | 0,377 | 0,709 | 0,73 | 0,571 |
| 8,952 | 8,838 | 6,787 | 7,727 | 6,877 | 8,025 | 8,016 |
| 0,252 | 0,257 | 0,255 | 0,234 | 0,374 | 0,37 | 0,306 |
| 0 | 0 | 0 | 0 | 0 | 0,001 | 0 |
| 0,013 | 0,013 | 0,003 | 0,004 | 0,006 | 0,007 | 0,008 |
| 0,002 | 0,002 | 0,002 | 0,002 | 0,002 | 0,002 | 0,002 |
| 0,002 | 0,002 | 0 | 0,001 | 0 | 0,002 | 0,002 |
| 1,31 | 1,3 | 1,998 | 1,929 | 1,734 | 1,728 | 1,837 |
| 2,397 | 2,847 | 1,853 | 1,595 | 1,839 | 1,705 | 1,731 |
| 2,894 | 3,034 | 1,172 | 1,119 | 1,234 | 1,271 | 1,331 |
| 7,199 | 6,904 | 0,684 | 0,793 | 2,088 | 1,522 | 1,362 |
| 0,468 | 0,355 | 1,488 | 0,879 | 1,559 | 1,457 | 1,213 |
| 0,041 | 0,046 | 0,033 | 0,028 | 0,04 | 0,035 | 0,04 |
| 0,324 | 0,342 | 0,147 | 0,138 | 0,151 | 0,158 | 0,156 |
| 0,576 | 0,585 | 0,266 | 0,275 | 0,274 | 0,283 | 0,309 |
| 0,634 | 0,664 | 0,405 | 0,369 | 0,503 | 0,492 | 0,45 |
| 1,184 | 1,257 | 0,987 | 3,695 | 0,462 | 0,419 | 0,723 |
| 0,411 | 0,388 | 0,032 | 0,212 | 0,448 | 0,26 | 0,423 |
| 0,006 | 0,006 | 0,005 | 0,005 | 0,006 | 0,01 | 0,008 |
| 0,09 | 0,088 | 0,041 | 0,166 | 0,038 | 0,07 | 0,094 |
| 0,552 | 0,545 | 0,121 | 1,02 | 0,124 | 0,554 | 0,399 |
| 0,033 | 0,033 | 0,035 | 0,036 | 0,04 | 0,053 | 0,049 |
| 29,218 | 30,089 | 24,433 | 19,096 | 24,05 | 21,356 | 23,748 |
| 0,01 | 0,01 | 0,011 | 0,01 | 0,012 | 0,015 | 0,012 |
| 0,057 | 0,055 | 0,018 | 0,051 | 0,053 | 0,059 | 0,05 |
| 0,006 | 0,006 | 0,007 | 0,01 | 0,006 | 0,013 | 0,01 |
| 0,015 | 0,002 | 0,002 | 0,001 | 0,002 | 0,005 | 0,007 |
| 0,005 | 0,006 | 0,005 | 0,005 | 0,006 | 0,008 | 0,006 |
| 0,03 | 0,019 | 0,033 | 0,023 | 0,031 | 0,034 | 0,038 |
| 0,006 | 0,005 | 0,006 | 0,005 | 0,007 | 0,008 | 0,006 |
| 0,01 | 0,009 | 0,007 | 0,007 | 0,009 | 0,01 | 0,009 |
| 0,005 | 0,005 | 0,001 | 0 | 0,004 | 0,002 | 0,002 |
| 0,169 | 0,183 | 0,019 | 0,016 | 0,044 | 0,039 | 0,061 |
| 1,045 | 0,909 | 0,14 | 0,054 | 0,42 | 0,168 | 0,313 |
| 0,011 | 0,003 | 0 | 0 | 0,013 | 0,001 | 0,001 |
| 0,032 | 0,03 | 0,008 | 0,008 | 0,014 | 0,014 | 0,016 |
| 0,039 | 0,037 | 0,032 | 0,034 | 0,04 | 0,043 | 0,041 |
| 0,001 | 0 | 0 | 0 | 0,001 | 0 | 0 |
| 0,01 | 0,007 | 0,002 | 0,001 | 0,004 | 0,004 | 0 |
| 0,248 | 0 | 0,036 | 0,017 | 0,122 | 0,06 | 0,086 |
| 0,028 | 0 | 0,005 | 0,001 | 0,012 | 0,002 | 0 |
| 0,015 | 0,012 | 0,001 | 0 | 0,01 | 0,003 | 0,007 |
| 2,503 | 3,068 | 1,879 | 1,874 | 2,996 | 2,828 | 3,058 |
| 0,005 | 0,004 | 0,004 | 0,004 | 0,005 | 0,007 | 0,006 |
| 0,009 | 0,011 | 0,007 | 0,01 | 0,008 | 0,009 | 0,007 |
| 0 | 0,038 | 0,011 | 0,042 | 0,027 | 0,011 | 0,025 |
| 0,005 | 0,004 | 0,052 | 0,053 | 0,034 | 0,03 | 0,001 |
| 0,453 | 0,472 | 0,588 | 0,575 | 0,678 | 0,761 | 0,647 |
| 0,008 | 0,01 | 0,01 | 0,01 | 0,009 | 0,009 | 0,009 |
| 0,004 | 0,004 | 0,007 | 0,008 | 0,006 | 0,006 | 0,004 |
| 0,005 | 0,006 | 0,005 | 0,005 | 0,007 | 0,007 | 0,006 |
| 0,09 | 0,105 | 0,123 | 0,123 | 0,123 | 0,143 | 0,137 |
| 2,943 | 3,292 | 3,464 | 3,565 | 4,31 | 4,444 | 4,536 |
| 0,079 | 0,046 | 0,065 | 0,124 | 0,096 | 0,084 | 0,025 |
| 0,138 | 0,143 | 0,158 | 0,168 | 0,179 | 0,209 | 0,202 |
| 0,001 | 0,002 | 0,004 | 0,004 | 0,003 | 0,003 | 0,004 |
| 0,015 | 0,015 | 0,018 | 0,014 | 0,016 | 0,016 | 0,002 |
| 0,026 | 0,026 | 0,029 | 0,026 | 0,031 | 0,036 | 0,03 |
| 0,13 | 0,127 | 0,144 | 0,146 | 0,161 | 0,181 | 0,16 |
| 0,239 | 0,248 | 0,275 | 0,265 | 0,298 | 0,332 | 0,311 |
| 0,076 | 0,06 | 0,093 | 0,101 | 0,076 | 0,079 | 0,057 |
| 0,122 | 0,122 | 0,146 | 0,145 | 0,16 | 0,181 | 0,17 |
| 0,082 | 0,086 | 0,099 | 0,1 | 0,105 | 0,121 | 0,114 |
| 0,019 | 0,019 | 0,022 | 0,021 | 0,024 | 0,026 | 0,025 |
| 0,09 | 0,082 | 0,105 | 0,092 | 0,09 | 0,112 | 0,108 |
| 3,076 | 3,246 | 3,224 | 3,065 | 3,764 | 4,042 | 3,514 |
| 0,501 | 0,517 | 0,351 | 0,357 | 0,374 | 0,415 | 0,417 |
| 0,003 | 0,003 | 0,004 | 0,004 | 0,003 | 0,004 | 0,004 |
| 0,003 | 0,003 | 0,004 | 0,003 | 0,002 | 0,005 | 0,005 |

| miR31_6 | miR31_7 | miR31_8 |
| --- | --- | --- |
| miR_31 | miR_31 | miR_31 |
| 0,505 | 0,489 | 0,372 |
| 6,815 | 8,588 | 9,084 |
| 0,316 | 0,308 | 0,236 |
| 0 | 0 | 0 |
| 0,002 | 0,001 | 0,001 |
| 0,001 | 0,002 | 0,001 |
| 0,001 | 0,002 | 0,001 |
| 1,636 | 1,711 | 1,708 |
| 1,399 | 1,112 | 1,124 |
| 0,949 | 0,707 | 0,776 |
| 0,108 | 0,104 | 0,489 |
| 2,946 | 3,347 | 2,893 |
| 0,023 | 0,027 | 0,032 |
| 0,087 | 0,086 | 0,097 |
| 0,217 | 0,125 | 0,183 |
| 0,411 | 0,332 | 0,326 |
| 2,713 | 1,656 | 5,665 |
| 0,134 | 0,161 | 0,311 |
| 0,006 | 0,008 | 0,008 |
| 0,128 | 0,167 | 0,158 |
| 0,828 | 0,814 | 0,937 |
| 0,039 | 0,046 | 0,043 |
| 28,225 | 19,812 | 20,874 |
| 0,011 | 0,012 | 0,011 |
| 0,049 | 0,043 | 0,026 |
| 0,009 | 0,013 | 0 |
| 0,002 | 0,015 | 0,01 |
| 0,004 | 0,006 | 0,006 |
| 0,023 | 0,034 | 0,035 |
| 0,005 | 0,005 | 0,001 |
| 0,005 | 0,005 | 0,006 |
| 0 | 0 | 0 |
| 0,053 | 0,003 | 0,015 |
| 0,031 | 0,07 | 0,075 |
| 0 | 0 | 0 |
| 0,005 | 0,006 | 0,009 |
| 0,035 | 0,039 | 0,039 |
| 0 | 0 | 0 |
| 0 | 0 | 0 |
| 0,008 | 0,015 | 0,021 |
| 0 | 0,001 | 0 |
| 0 | 0,001 | 0 |
| 3,014 | 2,553 | 2,135 |
| 0,003 | 0,005 | 0,005 |
| 0,004 | 0,01 | 0,008 |
| 0,024 | 0,008 | 0,046 |
| 0,048 | 0,067 | 0,063 |
| 0,608 | 0,649 | 0,578 |
| 0,007 | 0,009 | 0,009 |
| 0,005 | 0,006 | 0,007 |
| 0,006 | 0,006 | 0,006 |
| 0,103 | 0,12 | 0,131 |
| 3,875 | 4,525 | 3,896 |
| 0,105 | 0,139 | 0,013 |
| 0,15 | 0,201 | 0,196 |
| 0,004 | 0,005 | 0,005 |
| 0,004 | 0,012 | 0,007 |
| 0,018 | 0,023 | 0,012 |
| 0,121 | 0,154 | 0,146 |
| 0,241 | 0,297 | 0,287 |
| 0,054 | 0,092 | 0,059 |
| 0,136 | 0,158 | 0,156 |
| 0,082 | 0,101 | 0,107 |
| 0,021 | 0,024 | 0,024 |
| 0,069 | 0,078 | 0,076 |
| 3,146 | 3,789 | 2,946 |
| 0,362 | 0,432 | 0,417 |
| 0,003 | 0,004 | 0,004 |
| 0,003 | 0,003 | 0,004 |

| Sample Group | CTR_1<br>CTRL | CTR_2<br>CTRL | CTR_3<br>CTRL | CTR_4<br>CTRL | MiR31_1<br>MiR31 | MiR31_2<br>MiR31 |
| --- | --- | --- | --- | --- | --- | --- |
| IL_1a | 3 | 24,79 | 3,38 | 24,12 | 40,16 | 48,11 |
| IL_1b | 2 | 4,59 | 2 | 2,71 | 5,45 | 6,65 |
| IL_6 | 1 | 1 | 1 | 1,18 | 11,5 | 13,78 |
| IL_15 | 9,76 | 10,32 | 5,6 | 12,1 | 13,21 | 12,35 |
| IL_17 | 10,11 | 26 | 8,47 | 20,1 | 26,21 | 26,14 |
| IL_21 | 6 | 6 | 6 | 6 | 36,3 | 89,38 |
| TGFβ_1 | 404,7 | 354,58 | 263,3 | 376,9 | 606,15 | 536,27 |
| CCL13 | 11,86 | 5,43 | 5 | 23,48 | 29,13 | 32,01 |
| CCL22 | 25,35 | 19,45 | 8 | 20,63 | 20,55 | 20,19 |
| GM_CSF | 10 | 15,15 | 10 | 18,63 | 28,15 | 25,99 |
| sICAM | 732,39 | 435,38 | 253 | 632,67 | 762,21 | 635,11 |
| sCD163 | 190 | 450,79 | 190 | 734,08 | 718,31 | 823,56 |
| CCL2 | 532,72 | 500,27 | 500,43 | 493,72 | 949,93 | 677,83 |
| IL_8 | 299,58 | 259,38 | 293,06 | 271,95 | 490,16 | 398,95 |
| VEGF | 967,65 | 847,78 | 861,78 | 820,09 | 1412,28 | 1383,05 |
| S100A8 | 6920,26 | 5466,24 | 5355,07 | 5187,29 | 4268,85 | 4444,38 |

| MiR31_3 | MiR31_4 | MiR31_CB83 | MiR31_CB83 | MiR31_CB83 | MiR31_CB839_4 |
| --- | --- | --- | --- | --- | --- |
| MiR31 | MiR31 | MiR31+CB83 | MiR31+CB83 | MiR31+CB83 | MiR31+CB839 |
| 41,79 | 37,78 | 26,22 | 29,38 | 29,34 | 26,58 |
| 6,23 | 5,69 | 4,47 | 4,04 | 4,25 | 4,53 |
| 11,89 | 7,1 | 16,4 | 10,06 | 18,36 | 13,82 |
| 12,14 | 11,78 | 10,83 | 8,39 | 10,02 | 8,89 |
| 24,06 | 20,67 | 20,82 | 14,06 | 26,69 | 14,88 |
| 42,33 | 5,96 | 91,07 |  | 211,07 | 75,63 |
| 442,77 | 420,99 | 350,73 | 271,98 | 408,15 | 288,24 |
| 30,67 | 24,71 | 24,41 | 13,54 | 27,49 | 16,05 |
| 17,72 | 20,34 | 16,71 | 9,36 | 11,24 | 8,28 |
| 20,03 | 21,73 | 16,41 | 12,62 | 18,29 | 10,6 |
| 722,59 | 693,67 | 362,75 | 253,34 | 358,9 | 410,02 |
| 694,53 | 548,27 | 463,72 | 339,81 | 442,12 | 190,5 |
| 632,47 | 706,98 | 518,88 | 487,27 | 524,39 | 512,35 |
| 368,05 | 409,79 | 468,36 | 469,68 | 526,25 | 513,22 |
| 1216,55 | 1550,75 | 1393,86 | 1493,45 | 1609,5 | 1477,45 |
| 5243,36 | 6068,5 | 7388,46 | 5131,08 | 8404,95 | 7645,54 |

| Sample Group | CTR_1<br>CTRL | CTR_2<br>CTRL | CTR_3<br>CTRL | CTR_4<br>CTRL | TNF-α_1<br>TNF-α | TNF-α_2<br>TNF-α |
| --- | --- | --- | --- | --- | --- | --- |
| IL_1a | 3,38 | 24,79 | 3,38 | 24,12 | 48,96 | 44,79 |
| IL_1b | 2,70 | 4,59 | 2,70 | 2,71 | 9,62 | 6,77 |
| IL_6 | 1,00 | 1,00 | 1,00 | 1,18 | 177,09 | 235,53 |
| IL_15 | 9,76 | 10,32 | 5,60 | 12,10 | 17,67 | 21,90 |
| IL_17 | 10,11 | 26,00 | 8,47 | 20,10 | 45,66 | 37,36 |
| IL_21 | 6,00 | 6,00 | 6,00 | 6,00 | 260,62 | 300,41 |
| TGFβ_1 | 404,70 | 354,58 | 263,30 | 376,90 | 527,21 | 596,66 |
| MIP_1b | 1,33 | 9,99 | 3,90 | 12,05 | 13,62 | 15,13 |
| CCL13 | 11,86 | 5,43 | 5,00 | 23,48 | 33,49 | 29,62 |
| CCL22 | 25,35 | 19,45 | 19,00 | 20,63 | 146,46 | 215,30 |
| IP_10 | 0,00 | 0,00 | 0,00 | 0,00 | 301,57 | 424,43 |
| GM-CSF | 11,00 | 15,15 | 11,00 | 18,63 | 39,67 | 45,08 |
| sICAM | 732,39 | 435,38 | 435,00 | 632,67 | 2107,86 | 2105,99 |
| sCD163 | 450,00 | 450,79 | 450,00 | 734,08 | 1085,80 | 1056,48 |
| IL_8 | 299,58 | 259,38 | 293,06 | 271,95 | 6412,30 | 7763,62 |
| VEGF | 967,65 | 847,78 | 861,78 | 820,09 | 2176,97 | 2321,03 |
| S100A8 | 6920,26 | 5466,24 | 5355,07 | 5187,29 | 9099,30 | 8704,13 |

| TNF-α_3 | TNF-α_4 | TNF-α_CB83 | TNF-α_CB83 | TNF-α_CB83 | TNF-α_CB839_4 |
| --- | --- | --- | --- | --- | --- |
| TNF-α | TNF-α | TNF-α+CB83 | TNF-α+CB83 | TNF-α+CB83 | TNF-α+CB839 |
| 45,05 | 19,07 | 38,35 | 44,79 | 38,95 | 31,81 |
| 9,45 | 2,70 | 4,19 | 2,70 | 6,17 | 5,39 |
| 198,38 | 149,90 | 211,41 | 232,71 | 230,61 | 204,87 |
| 17,07 | 12,75 | 12,99 | 13,81 | 16,59 | 18,77 |
| 39,73 | 23,22 | 32,69 | 32,55 | 31,47 | 37,22 |
| 257,92 | NA | 23,49 | 158,90 | 178,11 | 75,63 |
| 487,71 | 455,77 | 387,68 | 384,51 | 521,17 | 439,67 |
| 12,77 | 6,86 | 16,34 | 10,28 | 12,15 | 11,61 |
| 36,32 | 19,99 | 27,23 | 16,55 | 36,23 | 31,57 |
| 118,07 | 138,08 | 103,65 | 197,16 | 168,59 | 192,19 |
| 319,51 | 296,86 | 421,06 | 617,80 | 553,67 | 570,05 |
| 33,45 | 11,80 | 20,05 | 21,75 | 24,14 | 21,63 |
| 1628,85 | 1623,70 | 2051,20 | 1731,99 | 2104,12 | 2071,17 |
| 812,00 | 330,64 | 690,56 | 610,02 | 678,60 | 741,94 |
| 6024,85 | 7050,43 | 3954,45 | 5409,96 | 4538,78 | 5072,56 |
| 1953,37 | 2173,86 | 2142,93 | 2573,08 | 2544,00 | 2599,08 |
| 8103,22 | 9490,44 | 6656,89 | 9050,13 | 9393,02 | 9636,13 |

**Extended Table 5:**
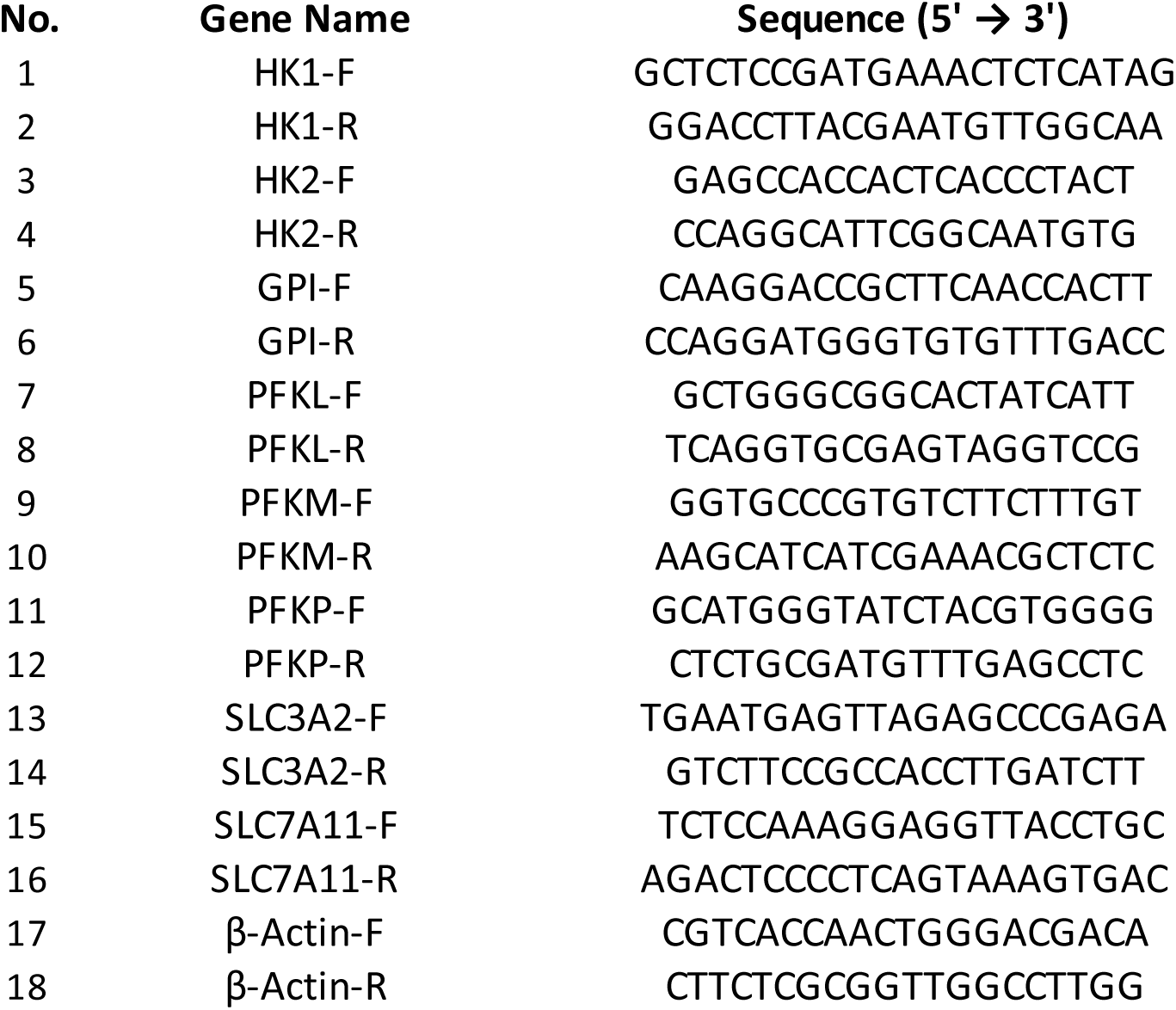
cytokine production in response to TNF-alpha and CB-839 treatment. (belonging to Luminex data, Fig. 4f)

**Extended Table 6:**
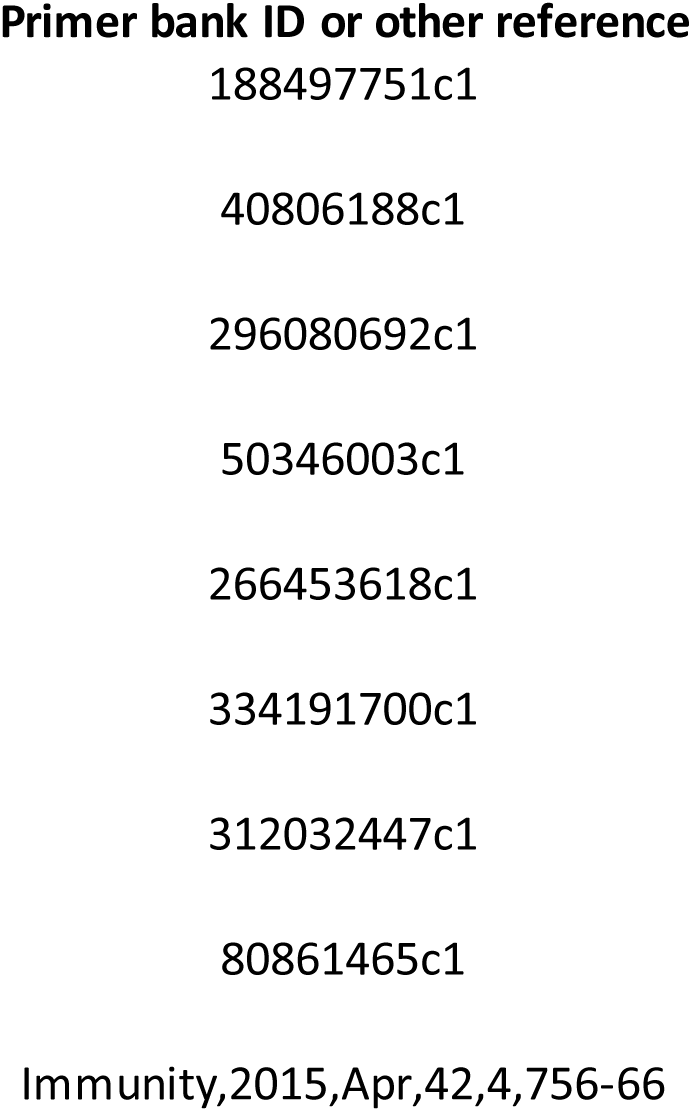
list of primers used for RTqPCR

**Extended Table 7:**
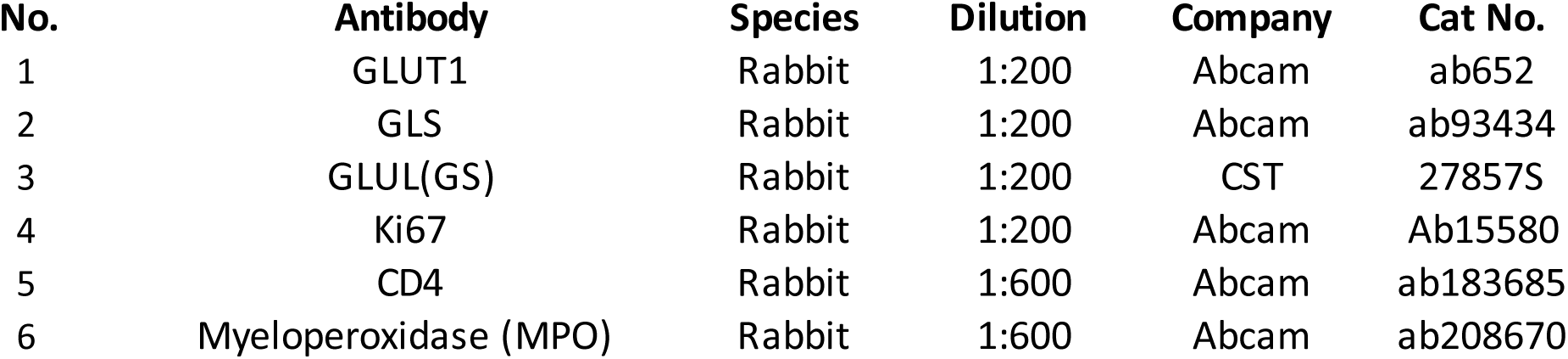
list of antibodies used for IHC study.

